# Diverse cell stimulation kinetics identify predictive signal transduction models

**DOI:** 10.1101/2020.01.28.923755

**Authors:** Hossein Jashnsaz, Zachary R Fox, Jason Hughes, Guoliang Li, Brian Munsky, Gregor Neuert

**Affiliations:** Department of Molecular Physiology and Biophysics, School of Medicine, Vanderbilt University, Nashville, TN 37232; Keck Scholars, School of Biomedical Engineering, Colorado State University, Fort Collins, CO 80523; Institut Pasteur, 25-28 Rue du Docteur Roux, 75015 Paris, France; Department of Chemical and Biological Engineering, Colorado State University, Fort Collins, CO 80523; Department of Biomedical Engineering, School of Engineering, Vanderbilt University, Nashville, TN 37232; Department of Pharmacology, School of Medicine, Vanderbilt University, Nashville, TN 37232

**Keywords:** predictive modeling, signaling, quantitative, kinetic cell stimulation, systems biology, single-cell, yeast, dynamic systems, dynamic response

## Abstract

The drive to understand cell signaling responses to environmental, chemical and genetic perturbations has produced outstanding fits of computational models to increasingly intricate experiments, yet predicting quantitative responses for new biological conditions remains challenging. Overcoming this challenge depends not only on good models and detailed experimental data but perhaps more so on how well the two are integrated. Our quantitative, live single-cell fluorescence imaging datasets and computational framework to model generic signaling networks show how different changing environments (hereafter ‘kinetic stimulations’) probe and result in distinct pathway activation dynamics. Utilizing multiple diverse kinetic stimulations better constrains model parameters and enables predictions of signaling dynamics that would be impossible using traditional step-change stimulations. To demonstrate our approach’s generality, we use identified models to predict signaling dynamics in normal, mutated, and drug-treated conditions upon multitudes of kinetic stimulations and quantify which proteins and reaction rates are most sensitive to which extracellular stimulations.

## INTRODUCTION

One of the longest standing challenges of modeling in systems biology has been to make accurate quantitative predictions for cell signaling responses over time, upon genetic mutations, when subjected to variable drug concentrations, and under kinetic changes of environment (Danhof et al., 2007; Rowland et al., 2011; Sebolt-Leopold and English, 2006; Wendell Lim & Bruce Mayer, 2015; Yaeger and Corcoran, 2019; Zanetti-Domingues et al., 2018). Examples of environmental perturbation include changes in the hormone or neurotransmitter levels (Steiner et al., 1982), morphogens (Sorre et al., 2014), or extracellular stressors such as osmolarity (Mettetal et al., 2008; Mitchell et al., 2015). Such perturbations are shown to vary significantly over both time and space and can affect cell fate decisions (Harvey and Smith, 2009). Signaling pathways are common in all eukaryotes and play key roles in cellular responses and function under a diverse range of different stimulations (Hatzivassiliou et al., 2010). However, mutations in these pathways can alter signaling dynamics and can contribute to many human diseases (Cildir et al., 2016; Hanahan and Weinberg, 2000; Logan and Nusse, 2004; Mitchell and Hoffmann, 2019; Suarez-Lopez et al., 2018). Therefore, understanding and predicting signal transduction network behavior will be a critical step to identify hidden regulatory mechanisms, distinguish between proteins and reaction rates that are sensitive to kinetic stimulations, and to detect and treat abnormal regulation that occurs in a large number of human diseases (Cildir et al., 2016; Hanahan and Weinberg, 2000; Logan and Nusse, 2004; Mitchell and Hoffmann, 2019; Suarez-Lopez et al., 2018).

A key obstacle that prevents predictive cell signaling models is the gross mismatch between the preponderance of biological complexity and the sparsity of quantitative experimental data (Oltvai and Barabási, 2002). Specifically, cell signal transduction networks are notoriously complicated (Campbell et al., 1998; Hanahan and Weinberg, 2000), yet experimental analyses of their dynamics often capture only a few signaling proteins at only a few time points during cellular responses (Kingsmore, 2006; Mitchell and Hoffmann, 2019; Muzzey and van Oudenaarden, 2009; Sorre et al., 2014). As a result, most current mechanistic models of signal transduction pathways are too complex and poorly constrained to provide predictive power, while current data-driven models are often too simple to extend beyond the most basic aspects of biological reality (Csete and Doyle, 2002; Klipp et al., 2005; Muzzey et al., 2009; Schoeberl et al., 2002). To address the disparity between biological complexity and lack of richness in experimental data, the dominant paradigm has been to devise (usually more expensive) experiments with higher content (e.g., high-throughput sequencing or multiplexed single-cell imaging) in hope that big enough data will eventually fill the gap between mechanistic and predictive understanding (Deng et al., 2019). Unfortunately, very little consideration has been given to the possibility that data could be richer if the experimental perturbations are designed with diverse kinetic features instead of constant environments of different concentrations.

To date, most computational models have been fit to experiments at steady state in different environments (Hao and O’Shea, 2012) or under step-like perturbations from one baseline level to another (Klipp et al., 2005; Sinkoe et al., 2017) (Figure 1A). However, life evolved to thrive in the presence of gradual changes (Harvey and Smith, 2009; Sorre et al., 2014), and recent studies have demonstrated that many cells respond differently to inputs of equal magnitudes based upon specific variations in input kinetics, such as different temporal frequencies (Albeck et al., 2013; Ashall et al., 2009; Cai et al., 2008; Hao and O’Shea, 2012; Hersen et al., 2008; Mettetal et al., 2008; Wang et al., 2012) or different spatial gradients (Harvey and Smith, 2009). A few pioneering studies have even demonstrated that different kinetic stimulations can dramatically affect intracellular signaling dynamics to create distinct cell phenotypes (Averbukh et al., 2018; Cai et al., 2004; Heltberg et al., 2019; Mettetal et al., 2008; Mitchell et al., 2015; Muzzey and van Oudenaarden, 2009; Rahi et al., 2017; Shimizu et al., 2010; Sorre et al., 2014; Thiemicke et al., 2019; Zhang et al., 2019) (Figure 1A). The fact that different kinetics of the same environmental inputs create such different responses offer a new opportunity to generate a wider and richer range of signaling pathway dynamics, while using current experimental assays that measure only a small number of signaling proteins.

**Figure 1.**
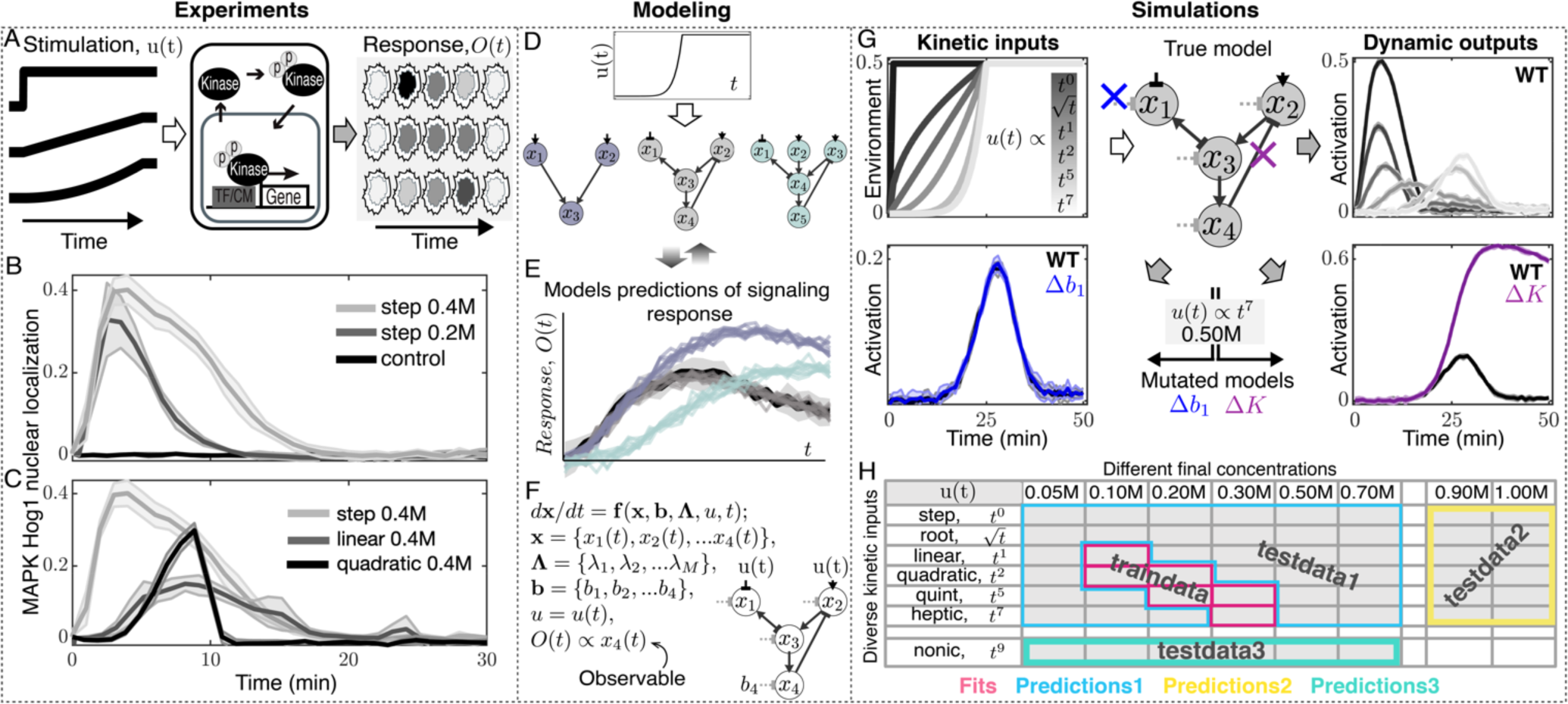
Kinetic stimulation of signaling pathways is required to identify predictive models and phenotypes. (A) Different extracellular kinetic stimulations (left) activate a signaling pathway (middle) and result in distinct dynamic kinase signaling and nuclear localization (right). (B) MAPK Hog1 nuclear localization dynamics upon step increases in NaCl to different final concentrations. (C) Hog1 dynamics upon step, linear or quadratic increases from 0M to 0.4M NaCl. Lines are means and shaded area are the standard deviation of multiple biological replicates (*STAR Methods*). (D) Schematic overview of signaling models of different complexity. The middle model is defined as the true model to simulate synthetic data. (E) Schematic overview of model identification based on models’ predictions. The true model was used to simulate synthetic data for pathway response (defined as the activation of node x_4_). Different models were fitted to the same data to identify parameters. These models were then used to make predictions for pathway activation, depicted as black solid line, upon a different kinetic input that result in different responses for different models (colored lines). (F) Pathways are modeled with ordinary differential equations (ODE), where nodes (e.g., x_1_-x_4_) represent a group of protein(s), top-layer nodes are regulated by the kinetic input (*u*(*t*)), basal regulators (e.g., b_1_-b_4_) act on each node, and the final-layer node (e.g., x_4_) represents the terminal signaling protein that is observed (Figure S1; *STAR Methods*). (G) Diverse kinetic perturbations corresponding to step (*t*^0^), root (√*t*), linear (*t*^1^), quadratic (*t*^2^), quints (*t*^5^), and heptic (*t*^7^) changes over time (left) are applied to the true model (middle) resulting in synthetic signaling activation dynamics (right). Blue and purple show simulated pathway activation from mutated models (**Δ**b_1_ and **Δ**K) under a heptic (*t*^7^) input function each compared to WT in black. In synthetic signaling data, lines are means and shaded area are the standard deviation of simulated replicates (*STAR Methods*). (H) Diverse kinetic inputs reaching different final concentrations result in 54 distinct simulated datasets for each WT or mutant model. These datasets are used to train models and test their predictions (Figures S1, S2, and S7).

The importance of utilizing multiple different dynamical inputs to characterize experimental processes has long been recognized in the concept of persistence of excitation in adaptive systems identification theory (Billings, 2013; Isermann and Münchhof, 2011; Nowak, 2002). This theory states that complex system (e.g., signaling pathways) mechanisms can be inferred from measurements of only a handful of coupled inputs (i.e., perturbations) and outputs (i.e., response measurements), provided that the inputs cover an appropriate set of orthogonal dynamics that excite the most important modes of the complex system. In this article, we explore the possibility that the diverse data sets achieved by cell signaling networks with multiple different inputs would provide such information and allow for more predictive models of complex biological pathways. With these models, we computationally screen and distinguish between proteins, reaction rates, and hidden regulatory mechanisms that are sensitive to kinetic environmental perturbations. To accomplish this task, we begin by inferring several models from single-cell time-lapse microscopy data for a well-known Mitogen-Activated Protein Kinase (MAPK) pathway. We then examine through simulation which types (or combinations of types) of experiments (kinetic inputs) are most likely to excite the appropriate system dynamics, to reveal well-constrained model parameters and mechanisms, and to enable quantitative predictions for new environmental or mutant circumstances. However, the modeling approach is generalizable to any other signal transduction pathway in any organism in healthy or diseased tissue.

To provide context for our exploration of optimal experiment design in the elucidation of cell signaling pathways, we choose the High Osmolarity Glycerol (HOG) MAPK signaling pathway (Figure 1A) in the yeast *Saccharomyces cerevisiae* (Saito and Posas, 2012; Tatebayashi et al., 2015). This eukaryotic model system includes all of the most relevant features of signaling networks, including a terminal signaling protein (Hog1) that is regulated through a *branched* upstream protein network that generally consists of kinases, phosphorelays, and sensors (Cuadrado and Nebreda, 2010; Laboucarié et al., 2017; Weston and Davis, 2007). These proteins can be regulated through phosphatases, autoregulation, feedback and feedforward loops that collectively modulate the robustness of response in the face of intracellular or extracellular perturbations. The Hog1 kinase is evolutionarily conserved from yeast to human (Hog1/p38) and its molecular components and interactions have been well characterized (Saito and Posas, 2012; Tatebayashi et al., 2015). Moreover, measuring Hog1 activation dynamics in single cells through its nuclear translocation dynamics using time-lapse epi-fluorescent microscopy allows for high temporal resolution measurements that are not possible with other methods (Mettetal et al., 2008; Muzzey et al., 2009). With this system, we have previously demonstrated experimentally that changing the extracellular osmolyte concentrations over time allows us to generate diverse kinetic input profiles that result in distinct Hog1 activation dynamics as pathway response output (Thiemicke et al., 2019) (Figures 1B and 1C). We now seek to explore what implications this richness of input-to-output dynamics can have on the possibility to identify signaling models, to distinguish between proteins, reaction rates, and hidden regulatory mechanisms that are sensitive to kinetic environmental perturbations, and to determine under what optimal experimental conditions can this identification be most reliably accomplished.

For clarity of exposition, we define “kinetic stimulation” as the extracellular environmental perturbation that is applied to cells as inputs, and “dynamic responses” as the resulting pathway activation outputs. We refer to the pair of “kinetic stimulation” and “dynamic response” as a signaling dataset. We design kinetic stimulations that change over time under a wide range of rates and intensities. Our analyses reveal that model predictability depends primarily on the diversity of kinetic cell stimulations rather than the amount or specific type of data used. We demonstrate the power of this approach in selecting the best predictive model and accurately predicting the loss and gain of function mutants’ responses such as feedback loops, phosphatase activity, and over and under expression of signaling proteins (Figure 1). By simulating data for a range of cell stimulations of different kinetic types and intensities (Figure 1; *STAR Methods*), we compare model fits (red) to different predictions (blue, yellow and green) and demonstrate that our findings are general, irrespective of the choice of kinetics that are used to evaluate the predictions.

## RESULTS

### Parametrizing signaling models with experimental data enables predictions of WT and mutant pathway responses upon kinetic stimulations

Since the HOG model pathway combines universal signaling network features as outlined above, it serves as a blueprint to build predictive signaling models of varying complexity (Figures 1A-1D). To begin, we parametrized several representative biologically inspired models (*STAR Methods*) by fitting them to experimental Hog1 nuclear localization data (Figures 1B-1D and S1K-M). Using experimentally constrained parameters for each model, we predicted Hog1 signaling dynamics upon different kinetic stimulation profiles. We chose the model whose least square errors were minimal in predicting experimental Hog1 dynamics (Figures 1D and 1E). The representative model in Figure 1D (middle), resembles a simplified *branched* signaling pathway consisting of four nodes, including one activating and one repressing sensor protein, constant basal regulators and a negative feedback loop from the terminal kinase to an upstream signaling branch (Figures 1F and S1). In the context of the HOG pathway, the node x_1_ represents the Sln1 branch including the proteins Sln1, Ypd1, Ssk1, Ssk2/ Ssk22. The Sln1 branch utilizes a two-component phosphorelay mechanism to transmit its signal with b_1_ representing the constant deactivation of the Sln1 branch (Hohmann et al., 2007; Maeda et al., 1994). A mutation in b_1_ could increase or decrease the activity of the Sln1 branch (Reiser et al., 2003). The node x_2_ describes the Sho1 branch of the HOG pathway that utilizes protein kinases to relay its information. The Sho1 branch consists of the proteins Sho1, Msb2, Hkr1, Opy2, Cdc42, Ste20/Cla4, Ste11 and Ste50 with b_2_ modeling the basal deactivation of the Sho1 branch (Tatebayashi et al., 2015). Changing or mutating b_2_ could result in an increase or decrease in the activity of the Sho1 branch. The Sho1 branch is also regulated by the Hog1 kinase through a feedback loop (Hao et al., 2007). Deletion of this interaction (**Δ**K) represents a Hog1 kinase dead mutant, a Hog1 analog sensitive mutant or an Hog1 inactivation due to a small molecule inhibitor (O’Rourke and Herskowitz, 1998; Westfall and Thorner, 2006). The node x_3_ represents a MAPKK such as Pbs2 that integrates information from two branches and has basal regulation (b_3_) through phosphatases such as Ptc1/2/3. Over or under expression of Ptc1/2/3 or mutations in Ptc1/2/3 that change phosphatase activity could then be modeled by changing the value of b_3_. Lastly, x_4_ represents a terminal kinase such as Hog1 that is activated by Pbs2. Hog1 is deactivated through phosphatases Ptc1/2/3 and Ptp2/3 (Mattison and Ota, 2000; Warmka et al., 2001; Young et al., 2002). Deactivation of Hog1 via constitutively active phosphatases is modeled as the act of basal deactivator b_4_ on x_4_. Changing the value of b_4_ in the model represents over- or under-expression of these phosphatases or a change in their activity. We defined the model in Figure 1D (middle) after parametrizing with experimental data as the “true model”. From this model we simulated synthetic signaling data upon diverse kinetic stimulations for the remainder of this study. Using a known model for this task, rather than additional experiments, allows us to systematically and quantitatively establish how diverse kinetic cell stimulations impact model identifiability and predictive power in a controlled setting where ground truth knowledge is available to check performance.

### Diverse kinetic cell stimulations result in distinct pathway activation dynamics

This known true model enables us to map diverse kinetic inputs to pathway dynamics in normal WT cells and mutant strains (Figure 1G). Over a wide range of kinetic input types and intensities, we simulated 54 synthetic datasets (Figures 1H, S2 and S7) under physiologically feasible and mutually independent kinetic stimulation profiles (see *STAR Methods*) such that each of the 54 profiles stimulates the pathway uniquely over time. Interestingly, the resulting pathway activation responses under different kinetic cell stimulations are qualitatively and quantitatively distinct from one another. Our simulated data (Figure S2) qualitatively and quantitatively captures the main characteristics of Hog1 dynamics, including activation level, measurement error, delay in activation, time to reach maximum activation and prefect adaptation observed in experiments (Figures 1B and 1C). Each signaling dataset covers the same duration and sampling range and has the same amount of data. However, different datasets may provide different amounts of information to constrain model parameters, which could lead to more or less accurate predictions for other kinetic inputs.

### Lack of kinetic stimulation diversity limits model prediction power

To explore what effects different stimulations have on model predictive power, we fit the true model to simulated data with experimentally realistic noise (*STAR Methods*) simultaneously for two steps of 0.2M and 0.3M NaCl (red). We then predicted the remaining 54 datasets for steps as well as all other kinetics (blue) (Figure 2). We performed multiple independent fits, where each fit over all time points on average converged to within the standard deviation in the synthetic data (Figures S1G-S1J; *STAR Methods*). Comparison of the fits and predictions to the corresponding training and testing data shows that the quality of predictions are nearly as good as fits for the *same* type of kinetics (Figures 2A and 2J, blue) but become worse for *different* types of kinetics (Figures 2B-2F and 2G-2I, 2K, 2L). This raises the question if the lack of predictability is due to the lack of data or due to limited kinetic diversity in the training data.

**Figure 2.**
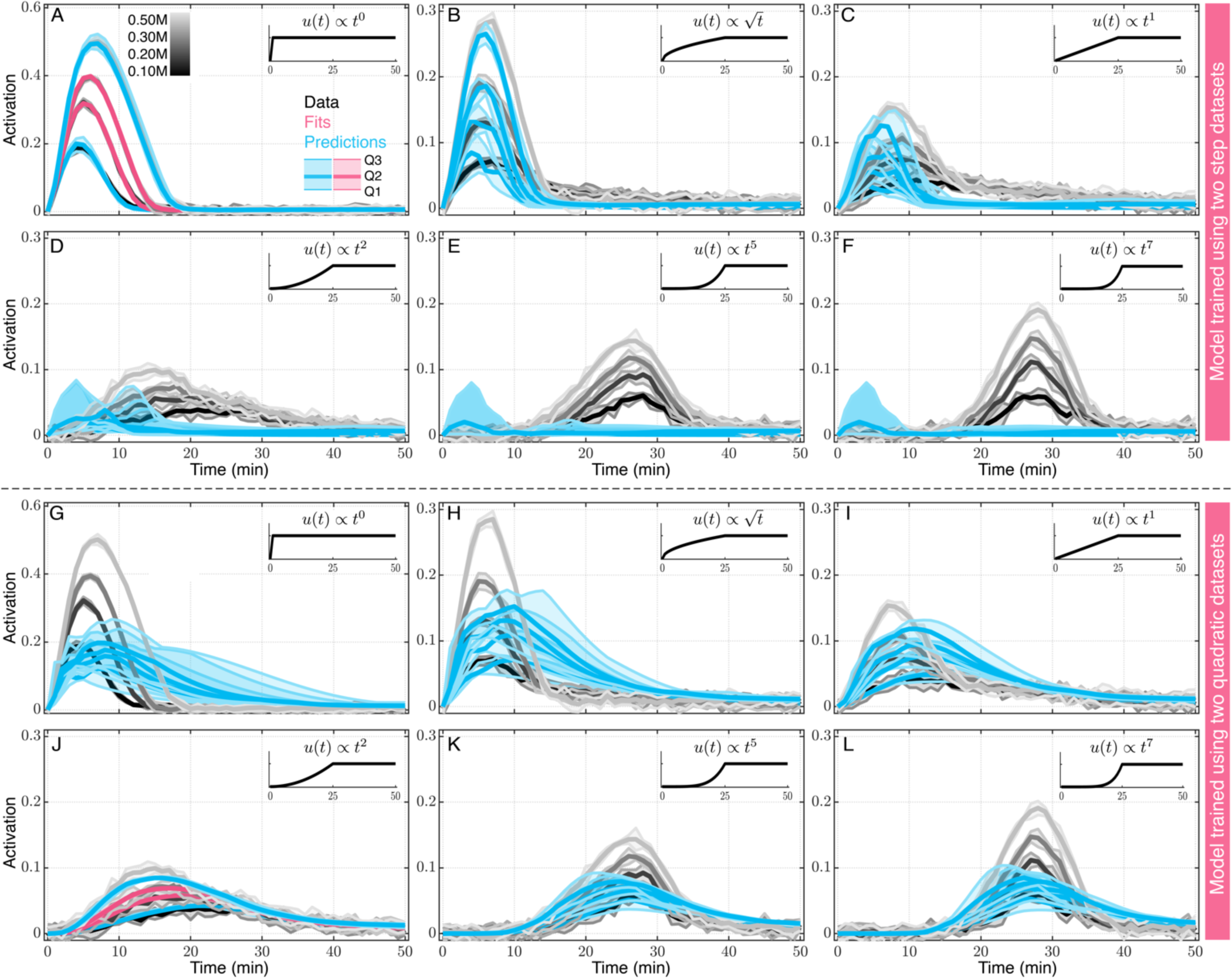
Models trained using same kinetic type inputs fail to predict pathway response to other kinetics. (A-F) Models trained using step inputs fail to predict pathway response to kinetic stimulations. Gray lines show synthetic pathway activation dynamics over time at different kinetic inputs as indicated inside each panel; step (*t*^0^, A), root (√*t*, B), linear (*t*^1^, C), quadratic (*t*^2^, D), quints (*t*^5^, E), and heptic (*t*^7^, F) input kinetics over time each to increasing final concentrations of 0.10M, 0.20M, 0.30M, and 0.50M. (A) Model fit simultaneously to steps of 0.2M and 0.3M data are shown in red. (A-F) Predictions under all other conditions are shown in blue. Predictions in (E) and (F) of all four concentrations overlap. Thick lines and shaded areas show median and interquartile range out of 10 independent fits and their corresponding predictions, respectively. As shown as an inset in (A), the 1^st^, 2^nd^ and 3^rd^ quartiles are used to plot shaded error bars where the thick line, upper and lower shaded areas represent Q2, (Q3-Q2), and (Q2-Q1), respectively. This convention is used throughout the manuscript. (G-L) similar to (A-F), models trained using quadratic inputs fail to predict pathway response to other kinetics.

To address the possibility of having too little data, we fit the model to six step (*t*^0^) data simultaneously (Figure 3A, red), and we predicted the signaling dynamics upon all remaining kinetic stimulations (48 data sets). We then compared model predictions of signaling dynamics for linear kinetic stimulations (*t*^1^) or nonlinear kinetic stimulations (*t*^9^) of different final concentrations to their corresponding synthetic data (Figure 3A, blue or green, respectively, compared to gray). After convergence, each set of model fits resulted in poor predictions for all data sets except for testing data collected using the same kinetic type as the training data. This observation was the same for subsequent training data sets with homogeneous input types (Figures S3A-S3C). Comparing how model predictability depends on the amount of training data of the same type illustrates that simply collecting *more* data of the same type does not automatically result in improved predictability (Figures 3B, S3A-S3C, right). Rather, the specific kinetics upon which the cells are stimulated may be of greater importance.

**Figure 3.**
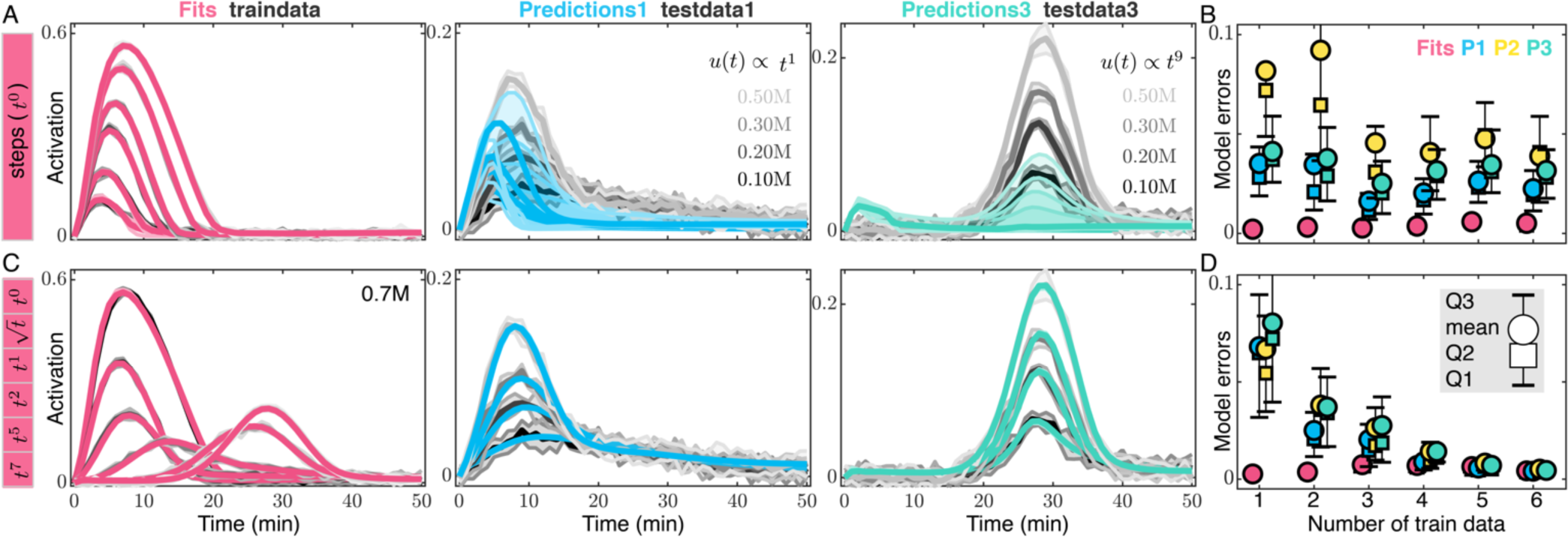
Kinetic stimulation improves model predictions. (A) Simultaneous fits (red) to six simulated step input response of different concentrations (gray) and subsequent model predictions of signaling dynamics upon different concentrations of linear input stimulations (predictions1 in blue, testdata1 in grey) or different concentrations of nonlinear inputs of the shape t^9^ (predictions3 in green, testdata3 in grey). (B) Box plots of mean and median fit and prediction errors when an increasing number of step inputs is used to train the model. (C) Simultaneous fit (red) to six different kinetic input stimulations of the same final concentration (gray) and model predictions for different concentrations of linear input stimulations (predictions1 in blue, testdata1 in grey) or different concentrations of nonlinear input stimulations of the shape t^9^ (predictions3 in green, testdata3 in grey). In (A and C), thick lines and shaded areas in gray show the mean and the standard deviation of synthetic data. Thick lines and shaded areas in red, blue, and green show median and interquartile range of 10 independent fits and their corresponding predictions, respectively. (D) Box plots of fit and prediction errors when an increasing number of diverse kinetics (*t*^0^ *to t*^7^) is used to train the model. For (B and D), circles, squares and error bars show mean, median and 1^st^ and 3^rd^ quartiles, respectively. Fit error statistics are drawn from n_train_ data sets over 10 independent fits (10 × n_train_ errors) where n_train_ = 1,2, …6 is the number of data sets used to train the model. Similarly, P1, P2, P3 are drawn from prediction errors of testdata1 (36-n_train_ data sets over 10 independent fits), testdata2 (12 data sets over 10 independent fits), and testdata3 (6 data sets over 10 independent fits), respectively (Figure 1H).

### Diversified kinetic stimulations better constrain model parameters and improve model predictions

To address the importance of kinetic diversity in training data, we fit signaling dynamics for different kinetic stimulations (*t*^0^ – *t*^7^) of a given final concentration (Figure 3C, red), and we predicted the signaling dynamics for the remaining kinetic stimulations (Figure 3C, blue and green). Comparing model predictions of signaling dynamics for linear kinetic stimulations (*t*^1^) or nonlinear kinetic stimulations (*t*^9^) of several different final concentrations to their corresponding synthetic data (Figures 3C and S3D, blue or green, respectively, compared to gray) indicates that all the predictions are substantially improved and are nearly as good as the fits, while the amount of training data is still the same as in Figure 3A. Quantitatively comparing how model predictability depends on the amount of different types of training data illustrates that kinetically diverse training data substantially improves predictability under all test kinetics (Figure 3D). A detailed analysis demonstrates that when using training data restricted to a single type, prediction errors increase as test data deviates away from kinetically similar training data (Figures 4A and S4A-S4I). To understand how diverse kinetic cell stimulations result in better predictions and reduced predictions uncertainties (Figure 4B), we developed a Fisher Information Matrix (FIM) analysis framework to directly estimate the uncertainties of model parameters under different experiment designs (Apgar et al., 2010; Fox and Munsky, 2019; Hagen et al., 2013; Jetka et al., 2018; Komorowski et al., 2011) (*STAR Methods*). For example, each row in Figure S3 represents such an experiment design. Comparing the estimates of the model parameters’ uncertainty using D-Optimality (determinant of FIM matrix), we find that parameters are constrained substantially better (smaller ellipse) via sets of experiments with diverse kinetics (Figures 4C and 4D, green, blue) compared to sets of experiments with the same amount of data but with homogeneous kinetics types (Figures 4C, 4D, and 4SJ-M, black, red, orange). These results indicate that signaling models built based on one type of kinetic cell stimulations may be predictive under different intensities of that same kinetic, but they fail to predict pathway responses upon other types of kinetic stimulations. This is important to consider because most computational models to date are often parametrized with measurements performed under constant stimulation profiles. Therefore, the models may not provide enough insights into cellular response in realistic kinetic physiological conditions. We find that training the model simultaneously with diverse kinetics constrains parameters betters and improves predictions substantially.

**Figure 4.**
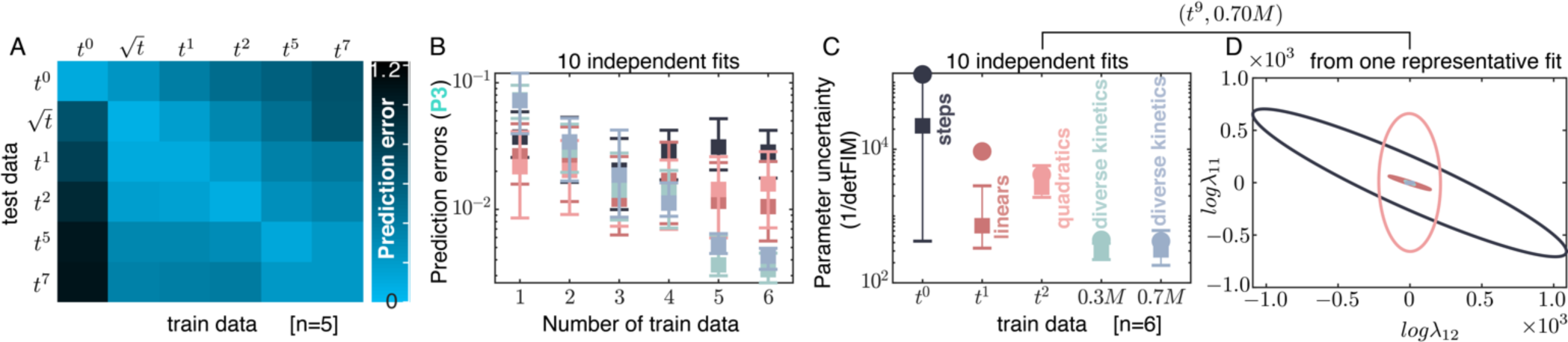
Kinetically diverse stimulations constrain model parameters substantially better than homogeneous kinetic types. (A) Comprehensive quantification of prediction errors of each kinetic stimulation type when five datasets of any given type is used to train the model (lighter colors denote smaller errors). (B) Comparison of prediction errors (Predictions3) under increasing amounts of training data of the same kinetic type (e.g., step, linear, quadratic) or diverse kinetic types (0.3M and 0.7M). (C) Parameter uncertainty of the model estimated as inverse of determinant of FIM (i.e., D-Optimality) when the model is fit to all six of each data set. (D) Ellipsis are representative 95% confidence intervals for a representative pair of parameters estimated from FIM^-1^. Colors correspond to the 5 different sets of experiments considered in C.

### Diverse kinetic stimulations improve model structure identification

Next, we examined how kinetically different cell stimulations affect model structure identifiability (e.g., the number and mechanisms of interacting signaling proteins) and elucidate the contribution of specific signaling proteins to overall dynamic signaling responses. Figures 5 and S5 show fits (red) and predictions (blue, yellow and green) of five models with varying complexity to six different signaling response dynamics. This analysis is performed using datasets that are simulated from model M3. Simpler models are built by removing one or two regulatory elements from M3 to form M2 or M1 respectively, to simulate two mutants of the true model where the corresponding kinase activities are removed resulting in loss of feedbacks regulations (Figures S5A-S5C). On the other side, a more complex model is built by adding an extra regulation element to the true model, which could represent a hidden regulatory element yet to be discovered (Figure S5D, model M4). Finally, another complex model is generated by adding an entirely new signaling branch (consisting of a third sensor node and introducing three additional regulatory elements) to the true model (Figure S5E, model M5). As expected, the simplest model cannot fit all data simultaneously (Figures 5A, S5A-S5B, red), whereas the true model and the more complex models fit well to the simulated data (Figures 5B, 5C, and S5C-S5E, red). These models were then used to make three sets of predictions (Figure 5D-5F, blue, green, yellow). As expected, the simple model does not predict well (Figure 5D), whereas the medium and complex models predict well but with varying levels of prediction uncertainties (Figures 5E, 5F). Through systematic comparison between different types of predictions for a range of concentrations and kinetic profiles (Figure 5G), we demonstrate that as model complexity increases, model fits improve as expected, but model predictability becomes worse as model complexity increases beyond the true model (Figure 5H). We argue that the increased uncertainty in the predictions of the complex models is due to the kinetically conditional behavior of their extra parameters and their flexibility in constraining their parameters during training the model (Figure S5). These results demonstrate that different kinetic cell stimulations can improve predictability in the process of complex model structure identification.

**Figure 5.**
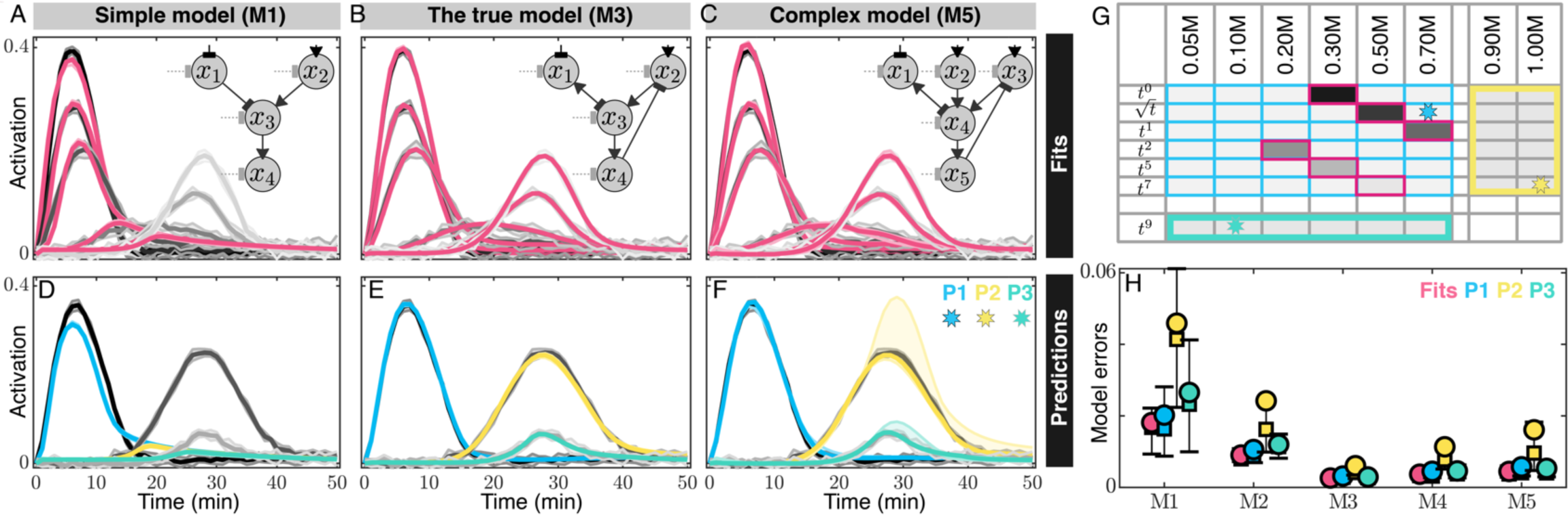
Kinetically diverse stimulation profiles enable unprecedented model predictions. (A-F) Predictions enable identification of the true model among models of increasing complexities. (A-C) Three models with increasing complexity from left to right (M1 in A, M3 in B, and M5 in C) each are trained with six kinetically diverse datasets that are simulated from M3. Model fits are shown in red and compared to training data in gray. (D-F) Model predictions (examples of Predictions1 in blue, Predictions2 in yellow, and Predictions3 in green) are compared to their corresponding test data (gray) indicated with stars in the table of train/test data in (G). Thick lines and shaded areas in gray show the means and the standard deviations of synthetic data. Thick lines and shaded areas in red, blue, yellow, and green show median and interquartile range of 5 independent fits and their corresponding predictions, respectively. (G) An overview of sets of training and testing data that are used for fits and predictions. Red, blue, yellow, and green squares indicate the data sets used in (H) and stars indicate predictions that are presented in (D-F). (H) Quantification of fit and prediction errors for five models of increasing complexity (see Figure S5 for model definitions and further analysis). Circles, squares, and error bars represent means, medians, and 1^st^ and 3^rd^ quartiles, respectively. Fit errors statistics are drawn from 6 train datasets over the 5 independent fits (30 fit errors). Similarly, P1, P2, and P3 are drawn from prediction errors of testdata1 (30 datasets, blue in G), testdata2 (12 datasets, yellow in G), and testdata3 (6 datasets, green in G), respectively, each collected over 5 independent fits.

### Diverse kinetic stimulations improve predictions of mutant responses

Finally, we examined how diverse kinetic stimulations affect predictive performance for *in silico* biologically realistic mutations to specific signaling proteins in signal transduction pathways (Figures 6A and S6) (Hohmann et al., 2007; Mattison and Ota, 2000; O’Rourke and Herskowitz, 1998; Westfall and Thorner, 2006; Young et al., 2002). Sensitivity analysis of WT model allowed to categorize model parameters into two main groups of insensitive (sensitivity = 0) and sensitive (sensitivity = 1) parameters that we then use to predict putative mutations (Figures 6A, 6B, and S6). To determine whether diverse kinetic cell stimulations can identify biological mechanisms, we computationally introduced six mutations in our true model (*STAR Methods*). These include three knockout mutants that remove basal deactivators on either x_1_ (**Δ**b_1_, blue cross), x_2_ (**Δ**b_2_, red cross), or x_3_ (**Δ**b_3_, teal cross). Mutants **Δ**b_1_ and **Δ**b_2_ can be interpreted as removing the constitutive deactivation of Sln1 or Sho1 branches, which could change the half-life of their active states. Mutant **Δ**b_3_ could represent regulation of Pbs2 through deletion of Ptc1/2/3 phosphatases. The node x_4_ can be mutated by removing kinase activity of x_4_ (**Δ**K, purple cross, e.g., kinase dead or kinase activity inhibited) and can be regulated through overexpression (*b*_4_, OE, orange); and underexpression (*b*_4_, UE, brown) of the basal deactivator on x_4_ such as the Ptp2/3 phosphatases (Figure 6A). We then simulated corresponding synthetic pathway activations from all six mutants upon all 54 kinetic stimulations (*STAR Methods*). One example for a simulation upon a t^9^ kinetic stimulation is shown in Figure 6C. To quantify the differences in signaling dynamics between normal and mutant cells, we define severity as the difference in the activation dynamics of a mutant compared to that of the WT (Figure 6D and *STAR Methods*). We observed that pathway activation in some mutant strains shows no difference from WT under all kinetic inputs (**Δ**b_1_ and **Δ**b_3_, mutation “severity” = 0), while other mutants with non-zero severity (**Δ**b_2_, **Δ**K, UE and OE) are different from WT (Figure 6C). Comparing the sensitivity to specific parameters in the model (Figure 6B) to the severity of the corresponding mutants’ effect on signaling dynamics (Figure 6E), highlights that sensitive parameters are direct indicators of how much specific mutations will affect signaling under different kinetics of specific types (Figure S6). Furthermore, constraining the parameters of WT model on its synthetic data under diverse kinetic stimulations enabled us to accurately predict the activation responses of all the 6 mutants over time under all 54 kinetic inputs tested (Figures 7 and S7). These results are quantitatively summarized in Figure 7E, showing that the prediction errors for simulated mutations are comparable to prediction errors of non-mutated WT cells.

**Figure 6.**
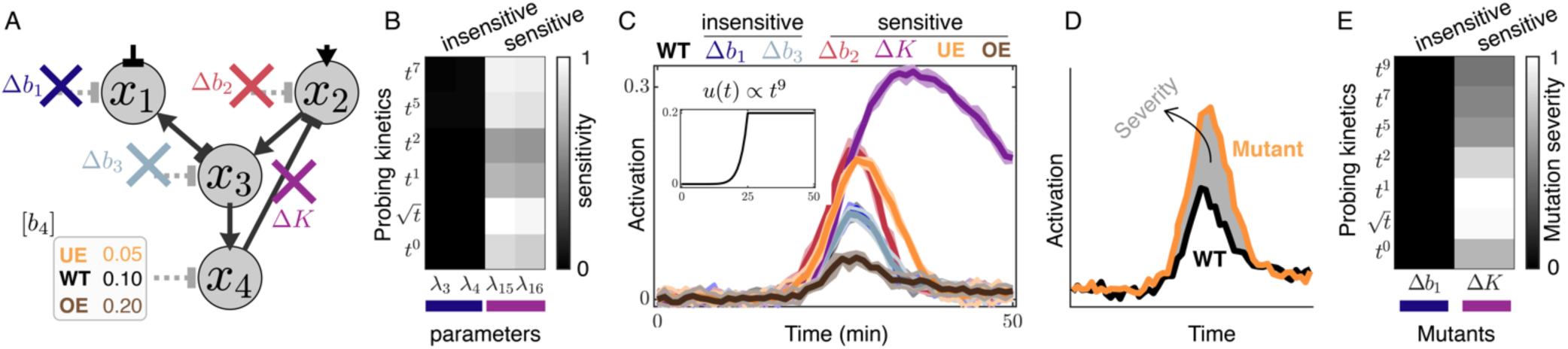
Kinetically diverse stimulations elucidate dynamic effects of mutants. Training the WT true model (M3 in Figure 5B) on its signaling dynamics upon diverse kinetics (red in Figures 5B and 5G) reveal insights into the response of several mutated models under all tested kinetic inputs. (A) The six mutants of the WT model correspond to deletions of basal regulators (e.g. phosphatases) on x_1_ (**Δ**b_1_, blue), x_2_ (**Δ**b_2_, red), or x_3_ (**Δ**b_3_, teal), removal of the kinase activity (e.g. kinase dead or inhibited MAPK) of x_4_ (**Δ**K, purple), and under- or over-expressing b_4_ (e.g. phosphatase) that regulates x_4_ (UE in orange and OE in brown). (B) Sensitivity analysis of WT model with respect to model parameters around their best value from the fits, predicts insensitive (e.g., *λ*_3_ and *λ*_4_ corresponding to **Δ**b_1_) and sensitive (*λ*_15_ and *λ*_16_ corresponding to **Δ**K) mutants. (See Figure S6). (C) Comparing the activation dynamics of mutants (simulated using Λ^0^ in Table S1) to WT under a representative kinetic input (t^9^ to 0.2M, inset). Thick line and shaded area (colors) show the mean and the standard deviation of synthetic data for the corresponding strains. (D) Mutation severity, defined as the difference in activation dynamics of a mutant from that of the WT (*STAR Methods*). (E) Mutation severity are shown for two representative mutant strains; **Δ**b_1_ (insensitive) and **Δ**K (sensitive) over all kinetic types summed over all their final concentrations.

**Figure 7.**
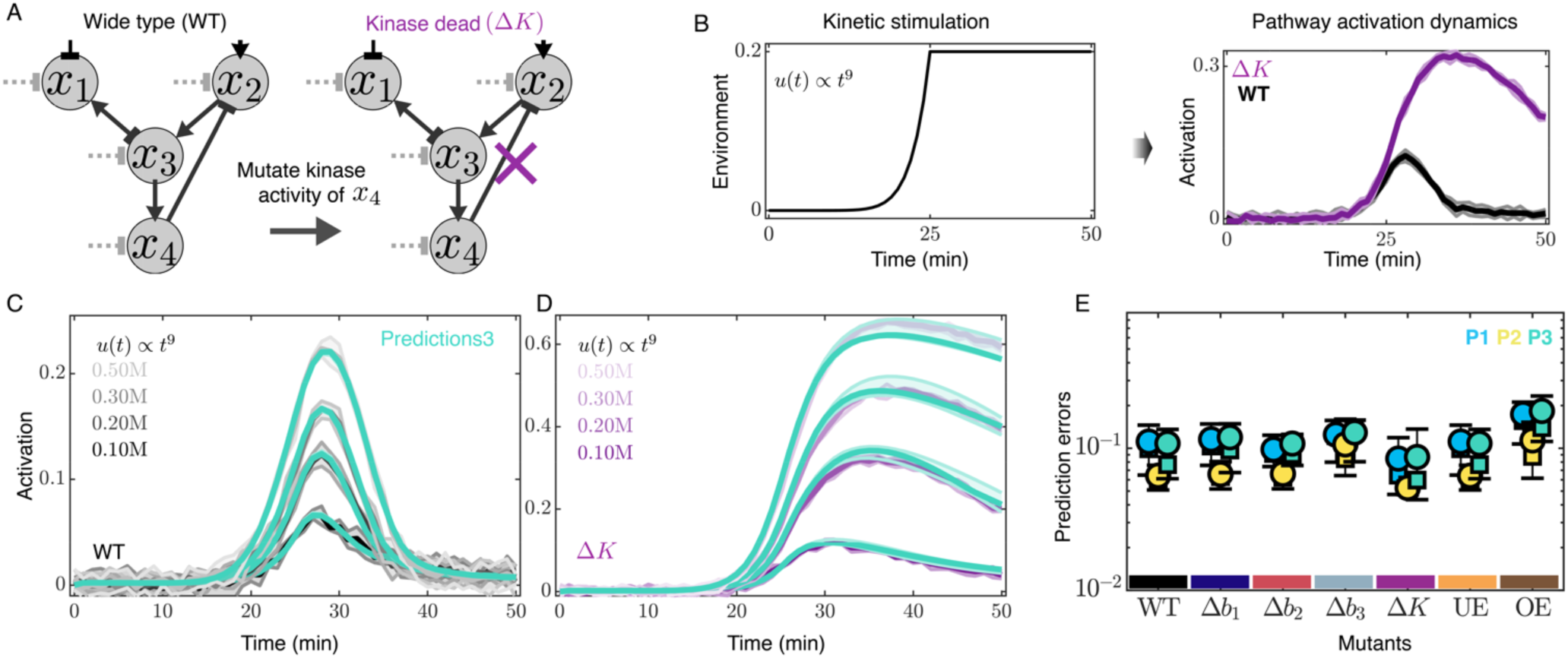
Kinetically diverse stimulations enable to predict mutants’ response dynamics. Training the WT true model (M3 in Figure 5A) on its signaling dynamics upon diverse kinetics enable predictions for the response of mutants upon all tested kinetic inputs. (A) A mutant where the kinase activity of x_4_ is eliminated (e.g., kinase dead or inhibited MAPK, purple cross) leads to a loss of feedback regulation from x_4_ on x_2_. (B) Extracellular stimulation of the models result in elongated response adaptation in Δ*K* mutant compared to WT. (C-D) Example pathway activation predictions (predictions3 in green) are compared to their corresponding synthetic data for WT (C, gray) and **Δ**K (D, purple) under representative *t*^’^ kinetic inputs. (E) Prediction (P1, P2, P3) errors quantified over all 54 kinetics (Figure 1H) for each of the six mutants (Figure 6A) compared to WT. These predictions are made using parameters constrained from 5 independent fits of WT model in Figure 5B.

## DISCUSSION

Our experimental and simulation results demonstrate that different kinetic cell stimulations of a pathway give rise to distinct signaling activation dynamics (Figure 1). When compared to the same amount of any type of homogeneous kinetics, kinetically diverse cell stimulations perform much better to constrain complex model parameter sets and result in significantly reduced predictions errors (Figures 3 and 4). The FIM analysis approach provides a rigorous and clear mathematical interpretation of this effect. Specifically, the eigenvector of the FIM corresponding to the greatest eigenvalue corresponds to the most accurately constrained parameter combination for a given kinetic stimulation (Figure S4M). By comparing eigenvectors corresponding to large FIM eigenvalues, it is easy to see which kinetic type may be most effective to constrain specific parameters of a complex regulatory network. Similarly, by examining eigenvectors corresponding to small FIM eigenvalues, it becomes obvious which parameter combinations cannot be precisely identified using a specific kinetic input. By choosing diverse and complementary input kinetics, such that the full parameter space is spanned by high-eigenvalue FIM eigenvectors from one or multiple kinetic inputs, it becomes possible to constrain the entire parameter set (Figures 4C, 4D, and S4J-S4M).

Better constrained parameters make it easier to identify predictive models of signal transduction (Figures 5 and S5). This also enables improved performance to predict pathway activation dynamics for protein mutant strains upon kinetic stimulations (Figures 6, 7, S6, and S7). To illustrate these predictions, we revisited the HOG pathway as an example (Figure 6). We predicted that mutations that alter b_1_ will not impact Hog1 signaling dynamics. Next we focused on the node x_2_ that describes the Sho1 branch in which mutating b_2_ could result in an increase or decrease in Hog1 signaling. We predicted that **Δ**b_2_ results in increased Hog1 signaling amplitude. In addition, the Sho1 branch can be altered in its activity by the feedback regulation from Hog1 kinase. In our model, we predicted that removing Hog1 kinase activity or inhibiting Hog1 kinase activity (**Δ**K) results in increased and prolonged Hog1 activation. Next, we focused on the node x_3_ which represents Pbs2. Our model predicted that alternations of phosphatases such as Ptc1/2/3 would not alter Hog1 signaling dynamics through changed Pbs2 activity (**Δ**b_3_). Lastly, x_4_ represents the terminal kinase Hog1 that can be deactivated through phosphatases Ptc1/2/3, and Ptp2/3 (b_4_). The model predicted that changing the value of b_4_ through over or under-expression has a strong impact on Hog1 signaling intensity. The experimental observations of the effect of above-mentioned mutants on the HOG pathway dynamics (O’Rourke and Herskowitz, 1998; Saito and Posas, 2012; Tatebayashi et al., 2015; Warmka et al., 2001) qualitatively validate our modeling results for all mutants (Figure 6). Using simulated data, we verified our mutant findings by quantitatively comparing predictions from each mutant to their corresponding synthetic data (Figure 7).

These results demonstrate that kinetic cell stimulations are ideally suited to discover novel regulatory interactions, reveal key functional proteins, identify predictive and biologically meaningful models, and help to gain novel biological insights. We believe that implementing diverse kinetic cell stimulations may provide new opportunities in constraining complex model parameters in situations where experimental design methods based on instantaneous changes in the environment such as steps or pulses of varying heights or frequencies have not provided great success (Billings, 2013; Isermann and Münchhof, 2011). Given the complexity of signal transduction networks (Campbell et al., 1998; Hanahan and Weinberg, 2000) and their limited response bandwidths (Hersen et al., 2008), steps or pulsatile stimulations of even varying intensities or frequencies may not provide enough kinetics to efficiently probe the rich dynamics underlying these networks (Billings, 2013; Steiner et al., 1982). Our approach on the other hand is widely applicable to many biological pathways that respond to a kinetic cell stimulation with a dynamic signaling response, and our approach has far reaching implications for predicting pathway response upon specific mutations or drugs (Campbell et al., 1998; Cuadrado and Nebreda, 2010; Hatzivassiliou et al., 2010). Being able to predict the pathway activation dynamics upon mutations or upon environmental changes may help design better drug treatments regimes. In addition, these results could benefit our understanding of human biology, particularly in areas such as optogenetics, gene regulatory networks, or synthetic biology, where predictive understanding of the system behavior with respect to extracellular kinetics or intercellular genetic perturbations are of immense interest (Aoki et al., 2019; Bashor et al., 2019; Csete and Doyle, 2002; Endy, 2005; Gardner, 2013; Harrigan et al., 2018).

## SUPPLEMENTAL INFORMATION

Supplemental Information includes Figures S1-S7 and Table S1.

## ACKNOWLEDGMENTS

GN is supported by NIH DP2 GM11484901, NIH R01GM115892 and Vanderbilt Startup Funds. JH is supported by NIH T32DK101003. ZRF and BEM are supported by NIH R35 GM124747. The authors thank Amanda Johnson, Alexander Thiemicke, Benjamin Kesler, Rohit Venkat, and Joseph Cleland for comments on the manuscript. This study used resources at the Advanced Computing Center for Research and Education (ACCRE) at Vanderbilt University, Nashville, TN (NIH S10 Shared Instrumentation Grant 1S10OD023680-01 (Meiler)).

## AUTHOR CONTRIBUTIONS

GN and HJ conceived the study. GN and BEM supervised the study. GN, BEM, HJ, and ZRF designed the study, and developed the modeling framework. HJ and ZF implemented the modeling framework, and HJ, ZRF, and JH performed the analysis. HJ, GL, and GN performed the experimental measurements and analysis of Hog1 dynamics. HJ and GN wrote the manuscript, and BEM, ZRF, and JH contributed in writing and editing the manuscript.

## STAR METHODS

### KEY RESOURCES TABLE

| REAGENT or RESOURCE | SOURCE | IDENTIFIER |
| --- | --- | --- |
| Software and Algorithms |  |  |
| MATLAB | Mathworks | www.mathworks.com |
| MATLAB code used for all simulations and calculations | this study | Available upon reasonable request |
| Experimental Models: Cell Lines |  |  |
| Yeast cells | Euroscarf | BY4741 |

### LEAD CONTACT AND MATERIALS AVAILABILITY

Further information and requests for resources should be directed to and will be fulfilled by the Lead Contacts, Brian Munsky and Gregor Neuert.

### METHOD DETAILS

#### Modeling pathway as a dynamic ODE system

A dynamic Ordinary Differential Equations (ODE) system is used to model the pathway as an enzymatic regulatory network (Figure S1). We developed a general framework that maps any arbitrary regulatory network to their corresponding ODE models implemented in MATLAB 2018a (Figures S1A-S1E). The framework dynamically takes arbitrary number of regulatory nodes in any topology and generates all possible ODE models corresponding to all the possible permutations of the regulations in the network. A convention used in Ref. (Ma et al., 2009) was adopted to formulate the rate equations of the model. Each node in our model represents a protein (or group of proteins) that has a fixed total concentration that can be interconverted between active and inactive states via regulations from either of the kinetic inputs, fixed basal regulators, or other nodes of the network (Figure S1A-S1C). All the regulations that a node receives are summed (Figure S1C), and each regulation (link) is modeled as a Michaelis-Menten function (Figure S1D). In the model, different numbers of nodes could act as sensors to receive extracellular stimulation and these regulators converge on a downstream node which in turn regulate a last node as the readout of the pathway. Each node receives a basal regulation from a source of a fixed concentration. This takes the opposite sign of the overall regulation the node receives from the input or internodes. Such regulation is considered for the role of constitutively active phosphatases and the autoregulation. Finally, feedforward/feedback loop (FFL/FBL) regulations are considered in the most general form in our model, such that complex dynamic behaviors like adaptation are possible. Any realization of such a network with basal regulations and internode regulations (including FFL or FFL regulation) can then be posed as a candidate model for fitting and predicting signaling data (see Figures 5 and S5 as an example).

Despite the apparent topological complexity of signal transduction networks, we focus on a general 4-node topology for the following reasons: i) it is well supported that there might only be a limited number of recurrent network topologies (“circuit motifs”) that are capable of robustly executing biological functions (Milo et al., 2002; Shen-Orr et al., 2002; Wagner, 2005). ii) Despite a large number of proteins involved in the signaling networks, multiple of these proteins can be grouped together and considered a virtual node without losing significant generality on the overall pathway activation dynamics and cellular response. Indeed, model reduction methods have been of interest to simplify complex biological systems by exploiting system properties such as time scales or parameters sensitivity analysis (Huang et al., 2010; Jeong et al., 2018). iii) Many signaling pathways are *branched* where two (or more) upstream multi-component branches (consisting of the sensors, their phosphorelays, or kinases) are receiving (either the same or different) stimulations through different mechanisms, then converge at a common component, which in turn regulate a terminal signaling protein. This class of *branched* pathways in their core could be most broadly modeled as a 4-node topology. For example, in the Hog1 MAPK signaling pathway in *S. cerevisiae,* either of the Sln1 or Sho1 branches could be grouped into one virtual node given their fast (millisecond) activation dynamics before they converge on Pbs2 compared to the longer activation dynamics of the Hog1 kinase (that is in the order of 5 minutes) (Saito and Posas, 2012; Tatebayashi et al., 2015).

#### Simulating synthetic pathway activation dynamics

To validate the modeling framework and more importantly to establish how model identification depends on the amount and the type of data, synthetic signaling activation dynamics was simulated from a known model in response to different kinetic stimulation profiles. The reason we simulate synthetic data for the part of this paper, in comparison to fully relying on experimentally measured data, is that synthetic data enables to explore how diverse kinetic cell stimulations impacts model identifiability and predictive power, without the obfuscation and potential unknowns that come from modeling experimental data. Challenges in using experimental data could come due to uncertainties in the model, the data, the integration of both, or simply undiscovered biology. Even in the rare cases where both model and measurements may be available, there is still a lack of understanding on how to integrate modeling frameworks with available experimental data such that meaningful new predictions could be made (Klipp et al., 2005; Muzzey et al., 2009). On the other hand, three main reasons make simulating synthetic data ideal for our purpose; i) To study how model predictions depend on data relies on availability of signaling dynamics over a wide range of kinetics and turning to synthetic data allows to simulate responses upon a wide range of perturbations. ii) Simulating data from a known model provides a ground truth to quantitatively benchmark the performance of a model identification framework while by using experimental data we don’t have an underlying known model to cross-check the results. iii) Similar to (ii) simulating synthetic data from a known model with known parameter values provides a reference point to fully parametrize the model. In addition, model performance could be tested in a wide range of parameter space. For these reasons we simulate signaling data under conditions that are biologically inspired and resemble experimental observations.

Among many models that equally fit and predict our Hog1 observation dynamics, a known network topology was chosen that could most broadly represent the class of ubiquitous *branched* signaling pathways and is parametrized through best fit to our available experimental Hog1 activation dynamics upon 0.2M and 0.4M NaCl (Figures S1A, and S1K-S1M). Using the resulting parameters (Table S1), synthetic pathway activation dynamics were generated that qualitatively and quantitatively recapitulate the experimental Hog1 pathway activation dynamics, such as activation levels, measurement noise, onset of activation, maximum activation time and perfect adaptation time (Figures S1 and S2). Upon each stimulation input, single cell trajectories with experimentally realistic noise (to capture cell to cell variability and measurement noise) were simulated (Figure S1E). We simulated 30 independent synthetic data sets for each condition under independent single-cell noise (Figure S2). We refer to these as 30 “synthetic replicates” of the same data that will be used to initiate 30 independent fits for each condition.

Data for a wider range of stimulations (20 different kinetics to 20 different final concentrations) was performed and conditions shown in Figure 1H were selected under the following criteria: i) to have a stimulation input profile that is physiologically feasible such that it could be generated and delivered to the cells in an experimental setup. Specifically, the solubility of the stimulus in the cell culture media and the operational rates range and precision of the syringe pumps determine the feasibility of generating a desired profile. All the cell stimulation profiles in this manuscript are physiologically feasible and experimentally achievable (Thiemicke et al., 2019). ii) To have a detectable pathway activation response (lower bound on the final concentration), and that the activation shows adaptation and does not saturate (higher bound on the final concentration). iii) To have mutually exclusive (independent) data so that not any pair are overlapping over time for both stimulation inputs and the corresponding pathway responses. This ensures each implemented profile input stimulates the pathway uniquely over time. These criteria guide the generation of biologically inspired synthetic data sets that enable the quantitative investigation of *whether* and *how* the type and the amount of data affect model predictions and model identifiability.

#### Fitting model to pathway activation data

We developed a customized optimization algorithm (implemented in MATLAB 2018a) to robustly and rigorously fit a given model to (any number of) pathway activation data to constrain the model parameters. One of the main challenges in parameter optimization is the quality and quantity of available experimental data (Figures 1 and S1). To deal with these limitations, we developed a combinatorial Genetic Algorithm (GA) that efficiently samples a large parameter space, combined with MATLAB’s built in routine *fminsearch* for finer tuning of the parameters at each minimum (Figure S1F). The algorithm dynamically takes a model and a set of training data (*D* = {*O*_1_(*t*), *O*_2_(*t*) … *O_d_*(*t*)}) and returns a set of parameter sets (**Λ**\*) that best fit the training data set (see *next section* in STAR Methods). For every condition presented, 30 independent fits were performed, each taking one of the 30 “synthetic replicates” of the simulated data along an independent random parameter initiation and resampling through the algorithm (Figure S1G-S1J). This ensures that results are statistically reproducible, that they are not artifacts of noise in the simulated data, and that they are independent of initial parameter guesses. All 30 fit optimizations converged as shown in Figure S1H (objective converges). Fit errors were calculated by comparing each of 30 fits to their corresponding “synthetic replicate” data. These errors were normalized with respect to the number of train data (*d*) and the number of time points in each data set.

#### Optimization algorithm

The algorithm is given in Figure S1F and runs for 21 iterations (*i* = {1,2, … 21}), which similar to other parameters used in the algorithm, is determined based on fits convergences. Each iteration goes through two Genetic Algorithm (GA) calls each followed by a fminsearch (light blue boxes). The first GA and its following fminsearch uses only 25% (selected randomly) of timepoints of each dataset in the train data (OBJ_25%_), which helps to escape the potential local minima. Then the second GA and its consequent fminsearch use all timepoints of the train data. At the first 4 iterations (*i* ≤ 4) as well as at every 4th (gold box), the 1^st^ GA takes N=200 parameter sets sampled uniformly in [-3,+3] in the logarithmic scale (base 10), and returns a parameter set (Λ^×^), which feeds into its following fminsearch and returns the parameter set Λ^G^. This parameter set is collected through the iterations, and it is used to resample the 200 parameter sets for the 2^nd^ GA, as well as for the 1^st^ GA for *i* > 4 that’s not every 4^th^ (purple boxes). Each GA runs for 20 generations, passes on 1 elite parameter set at each generation, and uses a custom mutation function that uses the best sets from the last generation (parents) to guess some new parameter sets. Objective values are updated during the 2^nd^ fminsearch if their value is improved. In total, 168,042 (= 2×21×20×200 + 2×21) number of parameter sets are evaluated for each model fit.

#### Predicting pathway activation dynamics

From the best parameter sets resulted from fitting the model to a training dataset, the model was solved for *x*_4_(*t*) (Figure S1E) to simulate the pathway activation prediction under any given stimulation kinetic input. All 30 independent predictions corresponding to 30 independent fits were computed using their corresponding best parameters sets (each of 30 **Λ**\*s, Figures S1I and S1J). Prediction errors were quantified by comparing each of 30 predictions to their corresponding “synthetic replicate” data. Prediction errors were normalized with respect to the number of time points.

#### Simulating synthetic data from mutant pathways

To evaluate the quality of the predictions upon different extracellular kinetic inputs in the presence of intracellular network perturbations, synthetic data was simulated under all kinetic stimulations from three main classes of mutations in the pathway for the true model, which are i) knockout mutations, ii) varying expression levels, and iii) inhibiting the activity of a protein (Figure 6A). Mutation data was simulated using the same set of parameters (**Λ**^0^, Table S1) with which the WT pathway activation data was simulated. Three different knockout mutations were generated by eliminating each of the basal regulators (*b*_1_, *b*_2_, and *b*_3_) acting on *x*_1_, *x*_2_, and *x*_3_ nodes. Knockouts are done be setting their corresponding parameters values in the model to zero. Another mutation where the activity of the last node (*x*_4_) (e.g., kinase dead) and thus its regulatory function in terms of feedback on the upstream node *x*_2_ was eliminated. This mutation was expected to show elongated perfect adaptation compared to WT. Finally, overexpression (OE) and underexpression (UE) mutations were generated by changing the concentration of the basal regulator (*b*_4_) acting on *x*_4_. Here *b*_4_ was set to 0.05 (in UE) and 0.20 (in OE) compared to *b*_4_ =0.10 of WT (a two-fold change for each). From each of the 6 mutated pathways of the true model, their corresponding activation dynamics, *O*(*t*) ∝ *x*_4_(*t*), was simulated under all the kinetic stimulation inputs (shown in Figure 1H), and representative activation dynamics for each mutant are shown in Figures S7A-S7F.

#### Predicting pathway activation dynamics for mutated pathways

For each of the 6 mutants, the best parameter sets (each of the 30 **Λ**\* resulting from each of the 30 independent fits) were used to generate their corresponding predictions under each extracellular kinetic input (Figures 6A and S7). For all results presented in this study on mutants, all **Λ**\*s are obtained by only fitting the WT pathway to its six dynamically different signaling responses; <u>no mutant models or data were used</u> (Figures 5B and 5G).

#### Fisher Information Matrix (FIM) analysis to estimate parameter uncertainties

The Fisher information matrix (FIM) analysis was used to estimate expected parameter uncertainty for different experiment designs (Apgar et al., 2010; Fox and Munsky, 2019; Hagen et al., 2013; Jetka et al., 2018; Komorowski et al., 2011). The FIM provides the amount of information an observable could provide around an unknown parameter, and it has been extensively used to estimate how well potential experiments will constrain model parameters

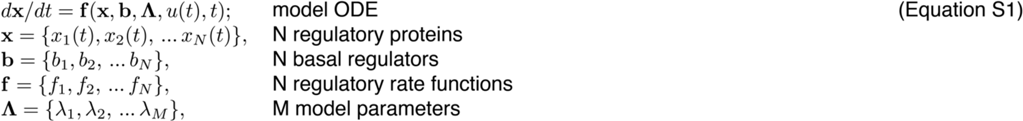

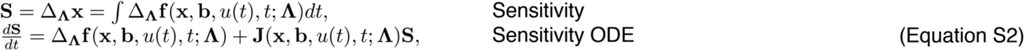

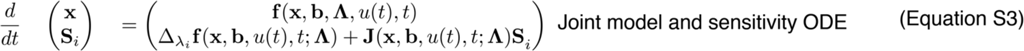

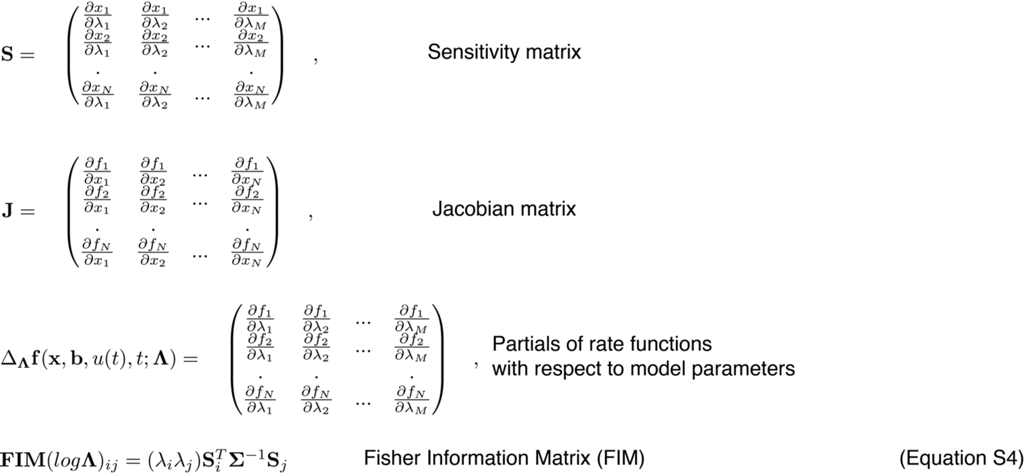

(Apgar et al., 2008; Bandara et al., 2009; Fox and Munsky, 2019; Sinkoe et al., 2017; Stewart-Ornstein et al., 2017). The FIM^-1^, the inverse of the FIM, known as the Cramer-Rao bound (CRB), is in particular useful as it provides a lower bound on the variance for any unbiased estimator of model parameters (Aitkin, 2010).

For any given model (Equation S1, Figure S1C), sensitivity equations for all model parameters (Equation S2) are formulated and are solved along the model ODEs (Equation S3) using Jacobina matrix of the rate functions under the initial and boundary conditions given in Figure S1E. Logarithmic parametrization of FIM is then computed to estimate the relative sensitivity of the parameters. From each model fit (to any set of train data), around each resulting best parameter set (**Λ**\*, Figure S1I) the FIM and its inverse, FIM^-1^, are computed upon all stimulation inputs. For each test data, standard deviation of the simulated activation dynamics over time, resulted from additive Gaussian noise that is independent of the mean activation (Figure S1E), is used to build the diagonal covariance matrix (Σ), which is used along computed sensitivities to calculate FIM (Equation S4). Several different metrics of the FIM, known as Optimality analysis, that are standard in model-guided experiment design are used to evaluate uncertainties (Fox and Munsky, 2019). These include A-Optimality, E-Optimality, T-Optimality, and D-Optimality, where the choice of the specific criteria depends on the application under the study. For example, E-Optimality corresponds to the smallest eigenvalue of the FIM, therefore gives a measure on how well an experiment design constrains the principle direction of parameter space that has the highest uncertainty. D-optimality, which corresponds to the determinant of the FIM provides a measure of the volume of the uncertainty in parameter space, therefore is best suited for our purpose to compare different experiments in their ability to constrain the model parameters. We defined a new optimality “W-Optimality” as a weighted sum over the uncertainties (Δ^*i*^) estimated by FIM^-1^ for individual parameters of the model (Figures S4J-S4M). In particular, this optimality gives more weight to better constrained parameters and less weight to insensitive ones, therefore it could provide a more accurate estimate on helpfulness of a specific experiment design in constraining the parameters that matters most by dynamically excluding the contribution of the sloppy parameters.

Under model fit to five different experiment designs that have the same amount of data (six steps, six linears, six quadratics, six diverse kinetics of 0.30M or six diverse kinetics of 0.70M), the FIM is calculated upon all test data. Then, the resulted FIM or FIM^-1^ was used to estimate the uncertainty of the model parameters using optimalities. Uncertainty estimated by each optimality is summed over all of testdata1, testdata2, or testdata3 kinetic stimulations and the results for 10 independent fits are shown in Figures 4C, 4D and S4J-S4M. The estimates of the model uncertainty for five different experiment designs computed using a representative kinetic input (t^9^, 0.7M) from 10 independent fits are shown in Figures 4C-4D for D-Optimality. A comprehensive analysis of the model uncertainty for all optimality criteria described above and using all kinetic stimulations are given in Figures S4J-S4M.

#### Mutation severity

For each mutant, mutation severity is computed as the sum of absolute difference in activation dynamics of a mutant from that of the WT over time upon all kinetic stimulations (Figure 6D). The mutation severity was marginalized for each kinetic type (summed over all final concentrations for each type kinetics), then normalized to the largest severity.

#### Experiments, image processing, and data analysis to measure Hog1 dynamics

Yeast *Saccharomyces cerevisiae* BY4741 was used for time-lapse microscopy. To assay the nuclear enrichment of Hog1 in single cells over time in response to NaCl osmotic stress, a yellow-fluorescent protein (YFP) was tagged to the C-terminus of endogenous Hog1 in BY4741 cells through homologous DNA recombination. A computer programmed syringe pump is used to control the osmatic stress over cells using a flowchamber^10^. The number of biological replicates (BR) and single cells presented in Figures 1B and 1C are as following; control has 6 BRs that have 6, 22, 5, 21, 9, and 9 cells; step 0.2M has 3 BRs that have 22, 53, and 67 cells; step 0.4M has 3 BRs that have 23, 25, and 42 cells; linear 0.4M 10min has 3 BRs that have 25, 9 and 45 single cells. quadratic 0.4M 10min has 3 BRs that have 67, 53 and 61 single cells.

### DATA AND CODE AVAILABILITY

The datasets of the current study is available from the corresponding author on reasonable request.

