## Supplementary_Information for "Diverse cell stimulation kinetics identify predictive signal transduction models"

**LIST OF SUPPLEMENTARY FIGURES**

**Figure S1. Signaling pathway, modeled as a dynamic ODE system, is parametrized using a custom optimization algorithm with Hog1 nuclear localization data.**

**Figure S2. Kinetic stimulations of signaling pathways result in dynamic pathway activation responses over a wide range of stimuli type and intensities.**

**Figure S3. Same kinetic type data fail to provide meaningful predictions regardless of amount of train data used.**

**Figure S4. Kinetically diverse stimulations constrain all model parameters while same kinetic type data are best effective on constraining some parameters but not others.**

**Figure S5. Upon diverse kinetics, predictions enable identification of a true model among models of increasing complexities.**

**Figure S6. Sensitivity analysis of parameters of the WT model trained upon diverse kinetics allows to screen for mutant responses.**

**Figure S7. Kinetically diverse stimuli enable to predict mutants' response dynamics.**

**LIST OF SUPPLEMENTARY TABLES**

**Table S1. Constrained parameter set that was used to simulate all the synthetic data in this paper from WT or mutant models.**

Figure S1

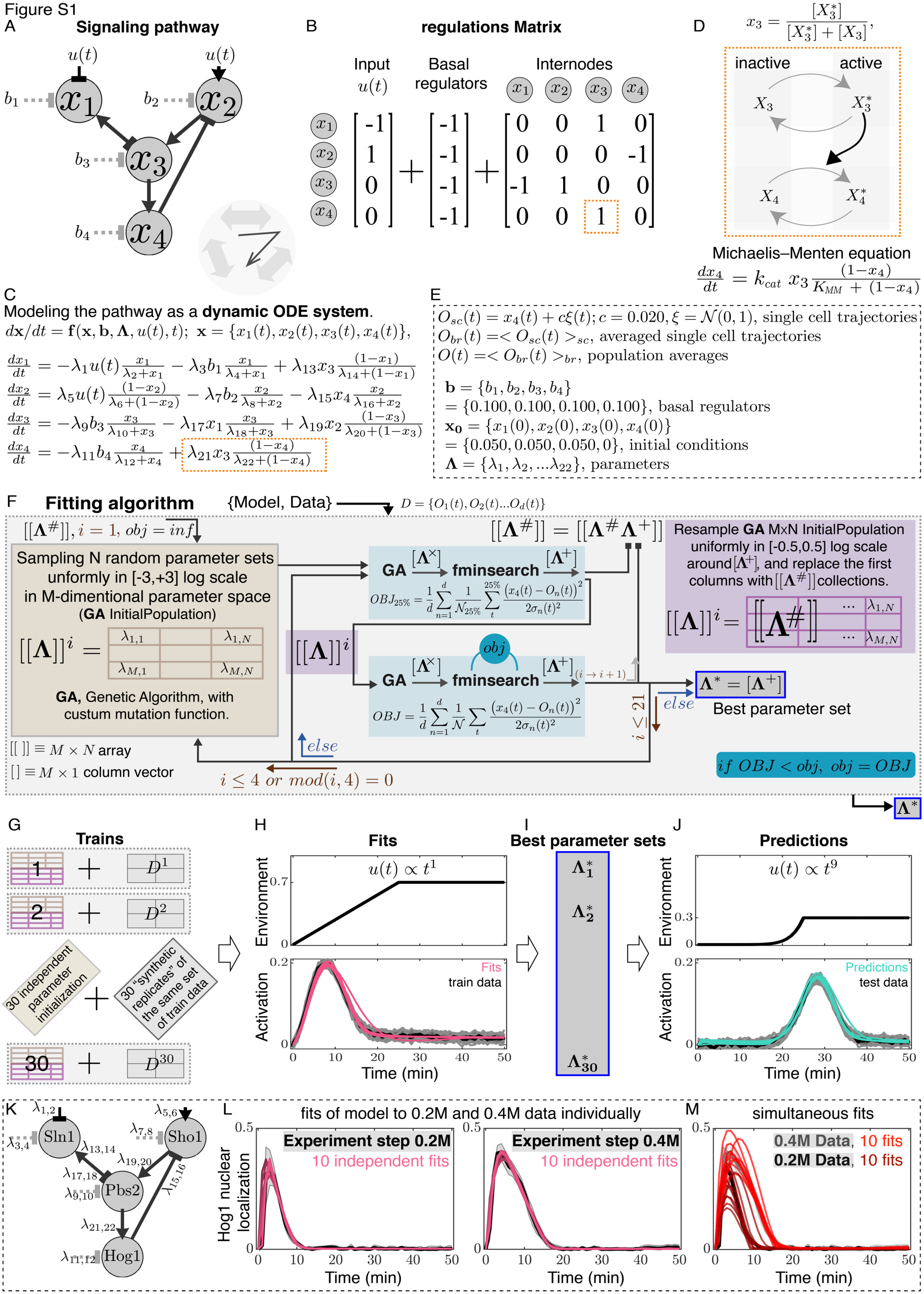
A
Signaling pathway
u(t)
u(t)
b1
b2
b3
b4
x1
x2
x3
x4
B
regulations Matrix
Input
u(t)
Basal
regulators
Internodes
C
Modeling the pathway as a dynamic ODE system.
$d\mathbf{x}/dt = \mathbf{f}(\mathbf{x}, \mathbf{b}, \mathbf{\Lambda}, u(t), t);\ \mathbf{x} = \{x_1(t), x_2(t), x_3(t), x_4(t)\},$
$\frac{dx_1}{dt} = -\lambda_1 u(t)\frac{x_1}{\lambda_2+x_1} - \lambda_3 b_1\frac{x_1}{\lambda_4+x_1} + \lambda_{13}x_3\frac{(1-x_1)}{\lambda_{14}+(1-x_1)}$
$\frac{dx_2}{dt} = \lambda_5 u(t)\frac{(1-x_2)}{\lambda_6+(1-x_2)} - \lambda_7 b_2\frac{x_2}{\lambda_8+x_2} - \lambda_{15}x_4\frac{x_2}{\lambda_{16}+x_2}$
$\frac{dx_3}{dt} = -\lambda_9 b_3\frac{x_3}{\lambda_{10}+x_3} - \lambda_{17}x_1\frac{x_3}{\lambda_{18}+x_3} + \lambda_{19}x_2\frac{(1-x_3)}{\lambda_{20}+(1-x_3)}$
$\frac{dx_4}{dt} = -\lambda_{11} b_4\frac{x_4}{\lambda_{12}+x_4} + \lambda_{21}x_3\frac{(1-x_4)}{\lambda_{22}+(1-x_4)}$
D
$x_3 = \frac{[X_3^*]}{[X_3^*]+[X_3]},$
inactive
active
Michaelis–Menten equation
$\frac{dx_4}{dt} = k_{cat}\ x_3\frac{(1-x_4)}{K_{MM} + (1-x_4)}$
E
$O_{sc}(t) = x_4(t) + c\xi(t); c = 0.020, \xi = \mathcal{N}(0,1)$, single cell trajectories
$O_{br}(t) =< O_{sc}(t) >_{sc}$, averaged single cell trajectories
$O(t) =< O_{br}(t) >_{br}$, population averages
$\mathbf{b} = \{b_1, b_2, b_3, b_4\}$
$= \{0.100, 0.100, 0.100, 0.100\}$, basal regulators
$\mathbf{x_0} = \{x_1(0), x_2(0), x_3(0), x_4(0)\}$
$= \{0.050, 0.050, 0.050, 0\}$, initial conditions
$\mathbf{\Lambda} = \{\lambda_1, \lambda_2, ...\lambda_{22}\}$, parameters
F
Fitting algorithm
{Model, Data}
$D = \{O_1(t), O_2(t)...O_d(t)\}$
$[[\mathbf{\Lambda}^{\#}]], i = 1, obj = inf$
Sampling N random parameter sets unformly in [-3,+3] log scale in M-dimentional parameter space (GA InitialPopulation)
GA, Genetic Algorithm, with custum mutation function.
$[[\ ]] \equiv M \times N$ array
$[\ ] \equiv M \times 1$ column vector
GA
fminsearch
$[[\mathbf{\Lambda}^{\#}]] = [[\mathbf{\Lambda}^{\#}\mathbf{\Lambda}^{+}]]$
Resample GA MxN InitialPopulation uniformly in [-0.5,0.5] log scale around $[\Lambda^+]$, and replace the first columns with $[[\Lambda^{\#}]]$ collections.
$\mathbf{\Lambda}^* = [\mathbf{\Lambda}^+]$
Best parameter set
else
$i \le 21$
$i \le 4\ or\ mod(i,4) = 0$
$if\ OBJ < obj,\ obj = OBJ$
G
Trains
30 independent parameter initialization
30 "synthetic" replicates of the same set of train data
H
Fits
$u(t) \propto t^1$
Environment
Activation
Fits
train data
Time (min)
I
Best parameter sets
J
Predictions
$u(t) \propto t^9$
Predictions
test data
K
Sln1
Sho1
Pbs2
Hog1
L
fits of model to 0.2M and 0.4M data individually
Experiment step 0.2M
10 independent fits
Experiment step 0.4M
10 independent fits
Hog1 nuclear localization
Time (min)
M
simultaneous fits
0.4M Data, 10 fits
0.2M Data, 10 fits

**Figure S1. Signaling pathway, modeled as a dynamic ODE system, is parametrized using a custom optimization algorithm with Hog1 nuclear localization data.** (A) The signaling pathway is represented as a network topology diagram and is modeled as an enzymatic regulatory network described by a dynamic ordinary differential equations (ODE) system represented in matrix (B) or equation (C) form. (D) A representative positive regulation in the model, where the active form of $x_3$ converts $x_4$ from its inactive to its active form (orange box in B and C) through a Michaelis-Menten function consisting of two parameters, the Michaelis-Menten constant ($K_{MM}$) and catalytic rate constant ($k_{cat}$). (E) The model is simulated with the initial and boundary conditions ($\boldsymbol{b}$ and $\boldsymbol{X_0}$) and Gaussian noise is added to the ODE solutions for $x_4(t)$ resulting in single cell pathway activation trajectories, $O_{sc}(t)$, under model parameters ($\boldsymbol{\Lambda^0}$) given in Table S1. Biological replicates are averaged from 10 simulated single cell trajectories. A signaling activation dynamic (e.g., each solid line and its shaded area in Figure S2) is calculated as the mean and the standard deviation of 5 simulated biological replicates. For convenience, we arrange the vector of parameters $\Lambda$ such that even entries correspond to MM constants and odd entries correspond to catalytic rate constants. (F) Optimization algorithm. A combinatorial optimization algorithm using a customized Genetic Algorithm (GA) combined with fminsearch. The algorithm takes a model and a set of train data, $D$, and returns a set of best parameter sets ($\boldsymbol{\Lambda}^*$) that best fit the train data. See STAR Methods. (G-J) Fitting-prediction method. (G) For every condition presented in the manuscript, 30 independent fits are performed, each taking one of the 30 “synthetic replicates” of the simulated data along an independent random parameter initiation and resampling through the algorithm shown in (F). Each dotted box in (G) is equivalent to (F). (H) From resulted 30 fits, either 10 or 5 best converged optimizations (as indicated in each figure in the manuscript) are kept (each red line), and from the (I) resulted best parameter sets, (J) their corresponding predictions (each green line) are made upon any kinetic stimulation input (STAR Methods). Here, the examples of fits (H, 10 red lines) and their corresponding predictions (J, 10 green lines) are shown from Figures 5A and 5C. In addition, at each best parameter set (I), upon each kinetic stimulation input, FIM (used to estimate parameter uncertainties) as well as sensitivity of the objective function with respect to parameters are also computed (STAR Methods). This method was used for all results throughout this manuscript. (K,L) Ten independent fits (reds) of model in (K) to measured Hog1 nuclear localization data (black) upon steps of 0.2M (left) and 0.4M (right) applied to the cells at 0 min. Solid black line is mean and shaded area the standard deviations out of multiple biological replicates (STAR Methods). (M) Ten best pairs out of 20 independent fits to 0.2M and 0.4M Hog1 data simultaneously (dark and light reds) of model in (K). Constrained parameters of the model (K) under this fit that result in best predictions are given in Table S1 ($\boldsymbol{\Lambda^0}$). This parameter set is used to simulate all synthetic data for the WT (Figure S2) and mutants (Figure S7) upon kinetic stimuli.

Figure S2

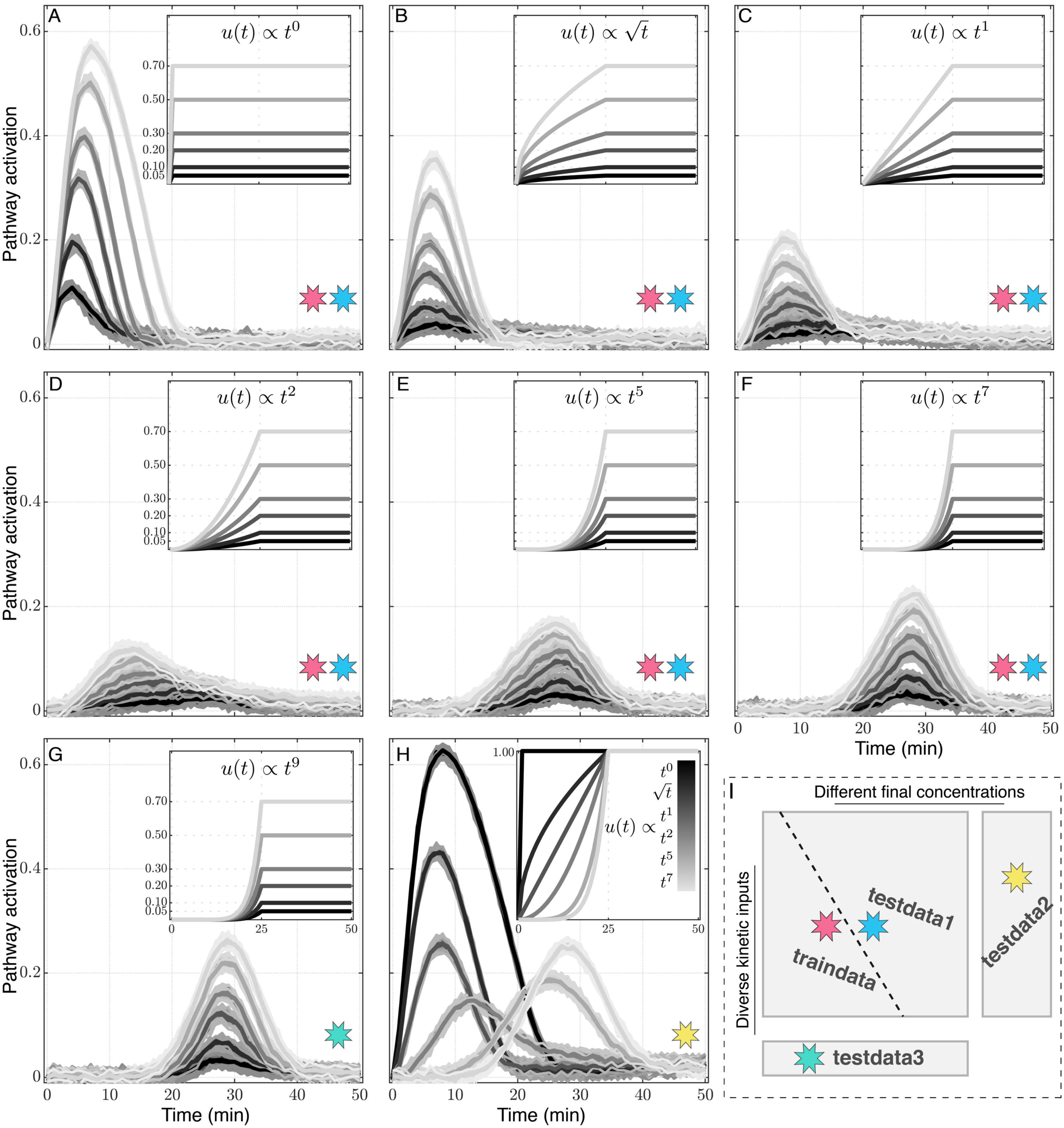

A
B
C
D
E
F
G
H
I
Pathway activation
Time (min)
$u(t) \propto t^0$
$u(t) \propto \sqrt{t}$
$u(t) \propto t^1$
$u(t) \propto t^2$
$u(t) \propto t^5$
$u(t) \propto t^7$
$u(t) \propto t^9$
$u(t) \propto$ $t^0$ $\sqrt{t}$ $t^1$ $t^2$ $t^5$ $t^7$
0.70
0.50
0.30
0.20
0.10
0.05
1.00
Different final concentrations
Diverse kinetic inputs
testdata1
testdata2
testdata3
traindata

**Figure S2. Kinetic stimulations of signaling pathways result in dynamic pathway activation responses over a wide range of stimuli type and intensities.** Synthetic pathway responses simulated from WT model shown in Figure S1A upon diverse kinetic stimulations (inserts) as step ($t^0$, A), root ($\sqrt{t}$, B), linear ($t^1$, C), quadratic ($t^2$, D), quints ($t^5$, E), heptic ($t^7$, F), and nonic ($t^9$, G) functions over time each to final concentrations of 0.050, 0.10, 0.20, 0.30, 0.50, 0.70M. All kinetic stimulations start at 0 min, all except steps reach their final concentrations at 25 min, then keep constant from 25 min to 50 min (insets). (H) Data from diverse kinetic inputs ($t^0$, $\sqrt{t}$, $t^1$, $t^2$, $t^5$, $t^7$) all to 1.00M final concentration. 30 "synthetic replicates" (STAR Methods) of each data is simulated under independent single cell noise (independent $\xi = \mathcal{N}(0,1)$ Gaussian noise). Solid black lines and shaded area represent the mean and the standard deviation of each data. (I) Red, blue, yellow, and green stars indicate each dataset belong to traindata, testdata1, testdata2, and testdata3, respectively (as in Figure 1H**),** to be used to train the model or to test model predictions.

Figure S3

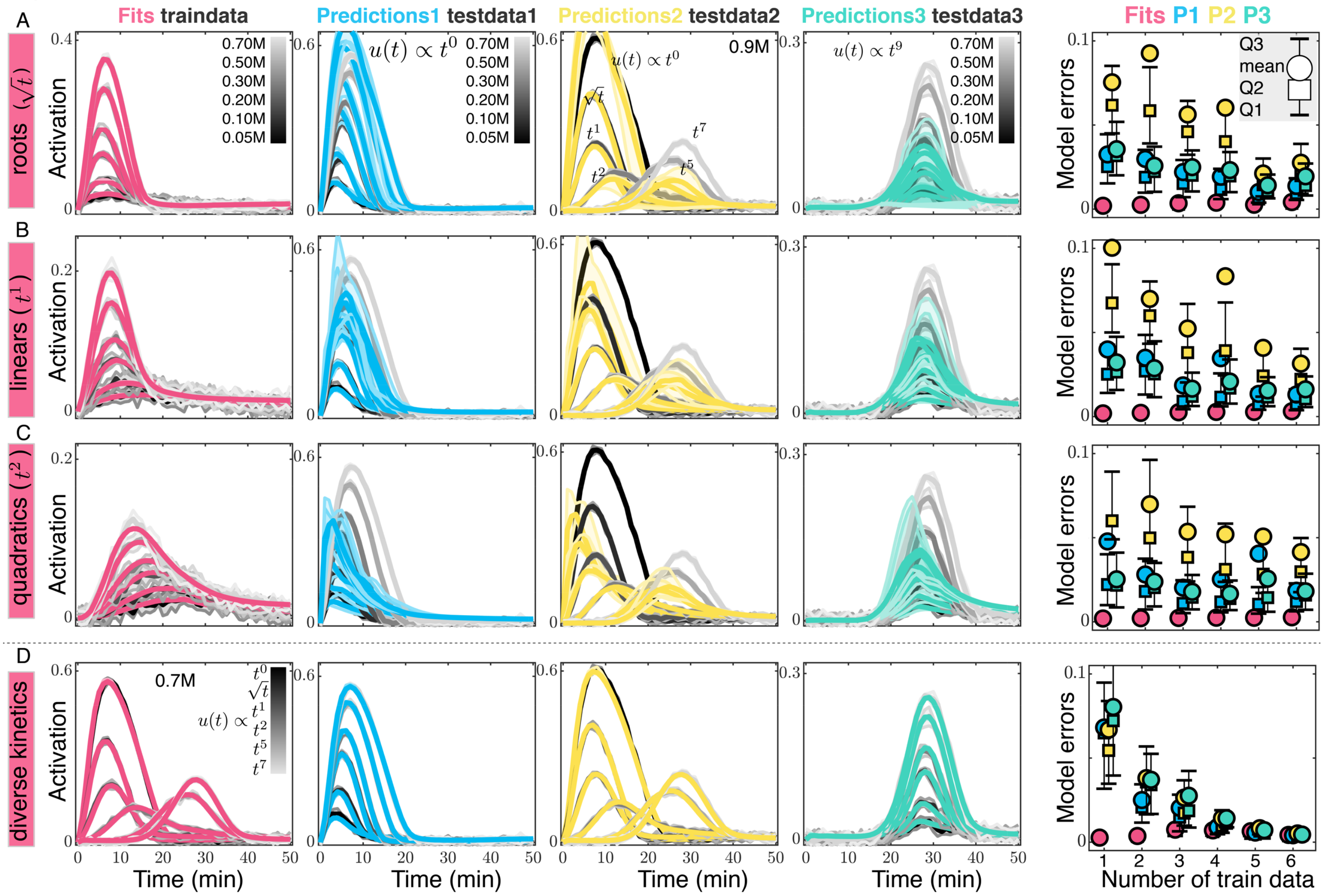
Fits traindata
Predictions1 testdata1
Predictions2 testdata2
Predictions3 testdata3
Fits P1 P2 P3
A
B
C
D
roots ($\sqrt{t}$)
linears ($t^1$)
quadratics ($t^2$)
diverse kinetics
Activation
Model errors
0.70M
0.50M
0.30M
0.20M
0.10M
0.05M
$u(t) \propto t^0$
0.9M
$u(t) \propto t^9$
$\sqrt{t}$
$t^1$
$t^2$
$t^5$
$t^7$
Q3
mean
Q2
Q1
0.7M
$u(t) \propto$
$t^0$
Time (min)
Number of train data

**Figure S3. Same kinetic type data fail to provide meaningful predictions regardless of amount of train data used.** (A) Model fits (red) simultaneously to all six root ($\sqrt{t}$) kinetics (gray), predictions (blue, yellow, and green) are compared to their corresponding test data (gray); Predictions1 (blue) to testdata1 (step kinetics, gray), Predictions2 (yellow) to testdata2 (0.9M diverse kinetics, gray), and Predictions3 (green) to testdata3 ($t^9$ kinetics, gray). Right: quantifications of fit and prediction errors when increasing number of root data are used to train the model. (B) Similar to (A) this time instead of all six root ($\sqrt{t}$) kinetics data, all six linear ($t^1$) kinetics data are simultaneously used to train the model. Fits (red) are compared to their corresponding train data (gray). (C) All six quadratic ($t^2$) kinetics data are simultaneously used to train the model. (D) Six diverse kinetics data are simultaneously used to train the model. In (B-D) predictions (blue, yellow, and green) are compared to their corresponding testdata in gray (similar to A). Thick line and shaded area in gray show the mean and the standard deviation of synthetic pathway activation data. Thick line and shaded area in red, blue, yellow, and greens show median and interquartile range of 10 independent fits and their corresponding predictions, respectively. Quantifications of model errors are similar to Figure 3 (upon all 54 kinetics over 10 independent model fits for each condition).

Figure S4

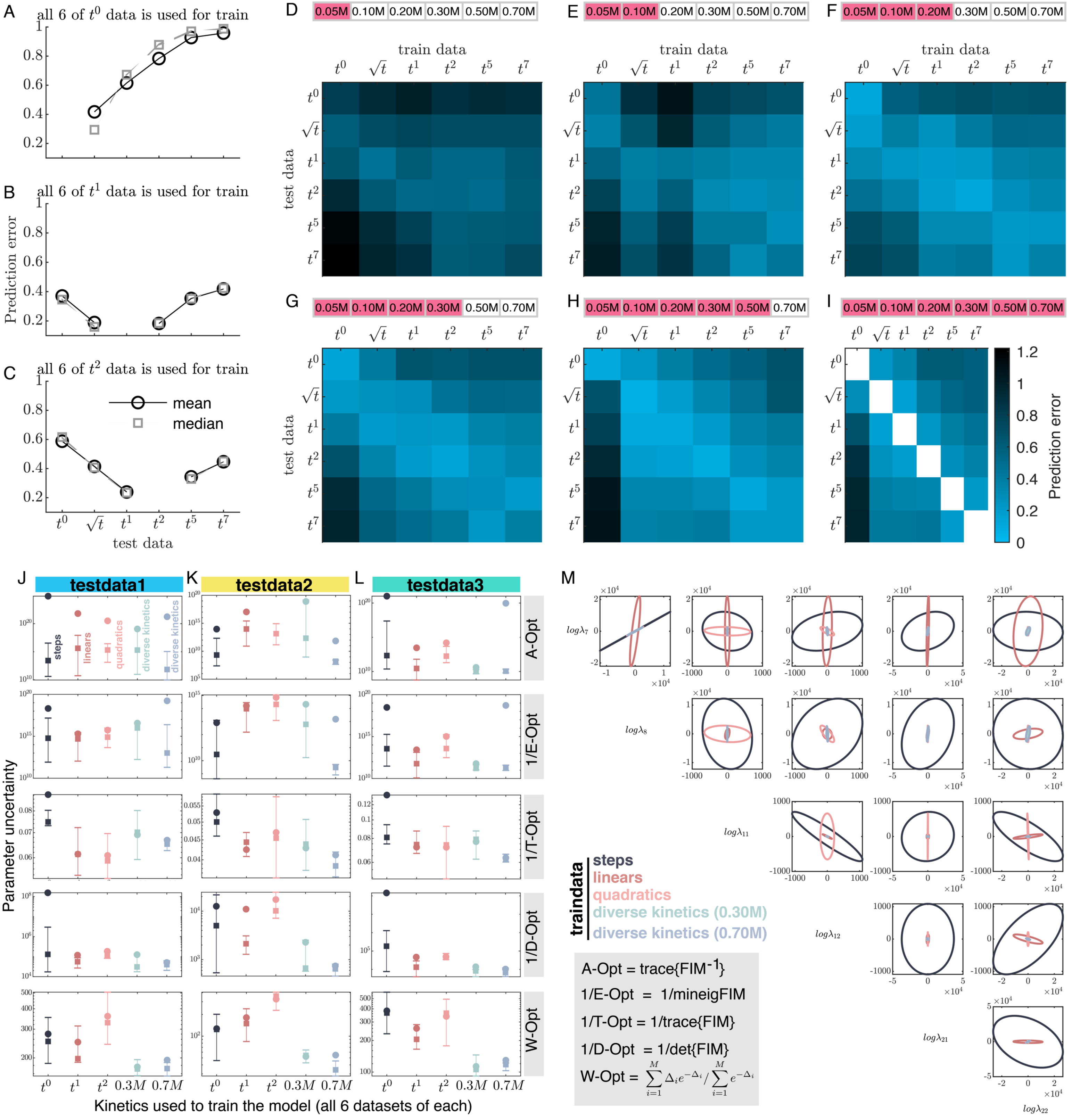
A
all 6 of $t^0$ data is used for train
B
all 6 of $t^1$ data is used for train
C
all 6 of $t^2$ data is used for train
Prediction error
mean
median
test data
D
E
F
G
H
I
0.05M 0.10M 0.20M 0.30M 0.50M 0.70M
train data
test data
Prediction error
J
testdata1
K
testdata2
L
testdata3
steps
linears
quadratics
diverse kinetics
A-Opt
1/E-Opt
1/T-Opt
1/D-Opt
W-Opt
Parameter uncertainty
Kinetics used to train the model (all 6 datasets of each)
M
$log\lambda_7$
$log\lambda_8$
$log\lambda_{11}$
$log\lambda_{12}$
$log\lambda_{21}$
$log\lambda_{22}$
traindata
steps
linears
quadratics
diverse kinetics (0.30M)
diverse kinetics (0.70M)
A-Opt = trace{FIM$^{-1}$}
1/E-Opt = 1/mineigFIM
1/T-Opt = 1/trace{FIM}
1/D-Opt = 1/det{FIM}
W-Opt = $\sum_{i=1}^{M} \Delta_i e^{-\Delta_i} / \sum_{i=1}^{M} e^{-\Delta_i}$

**Figure S4. Kinetically diverse stimulations constrain all model parameters while same kinetic type data are best effective on constraining some parameters but not others.** (A-I) Prediction errors increase by moving away from kinetically similar training data. (A-C) Quantification of prediction errors of each kinetic type when all six of steps (A), linears (B), or quadratics (C) are used to train the model. Kinetically similar data are best predictable, while by moving away from the kinetically similar train data predictions worsens quickly and dramatically. (D-I) Increasing the number of data (indicated with red boxes) from each type kinetics that is used to train the model (indicated on the x axis), and the predictions of each type of the kinetics (y axis) are quantified. Principal diagonals correspond to predicting test data that is kinetically similar to the train data (different final concentrations), which results in the best quality of predictions. All quantifications are from 10 independent fits for each condition. (J-L) Kinetically diverse stimulations constrain model parameters substantially better than same kinetic types. Parameter uncertainty (magnitude) estimated via different optimalities of FIM under different kinetic stimulations. Under five different sets of traindata including six steps, six linears, six quadratics, six diverse kinetics of 0.30M or six diverse kinetics of 0.70M, FIM and $FIM^{-1}$ are calculated upon all test data. The estimates of the model parameters uncertainty using each of A-Optimality, E-Optimality, T-Optimality, D-Optimality, and W-Optimality summed over for all of testdata1 (J), testdata2 (K), or testdata3 (L) kinetics and their boxplots out of 10 independent fits are shown for each condition. The fits are from Figures 3, S3 and S4I. W-Opt is defined as a weighted sum over the uncertainties ($\Delta^{i}$) estimated by $FIM^{-1}$ for individual parameters of the model (STAR Methods). (M) Ellipses as eigenvectors representing 95% confidence intervals estimated from $FIM^{-1}$ for few representative pairs of parameters. To clarify, similar results are obtained for all the model parameters. Under five different sets of traindata presented in J-L, as five different experiment designs, corresponding example ellipses are shown as indicated in the legend.

Figure S5

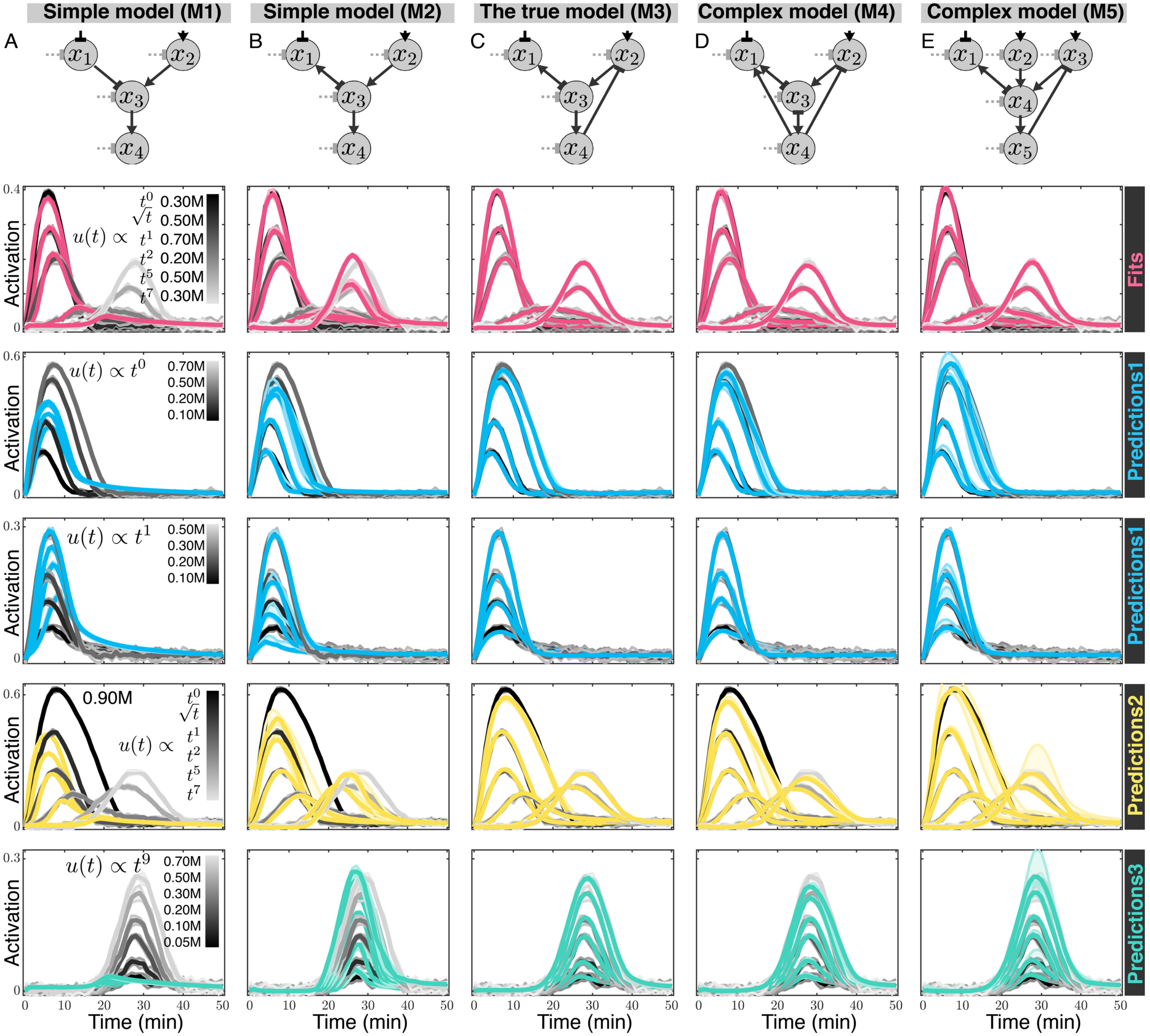
Simple model (M1)
Simple model (M2)
The true model (M3)
Complex model (M4)
Complex model (M5)
A
B
C
D
E
Activation
Time (min)
$u(t) \propto$
$t^0$ 0.30M
$\sqrt{t}$ 0.50M
$t^1$ 0.70M
$t^2$ 0.20M
$t^5$ 0.50M
$t^7$ 0.30M
$u(t) \propto t^0$
0.70M
0.50M
0.20M
0.10M
$u(t) \propto t^1$
0.50M
0.30M
0.20M
0.10M
0.90M
$u(t) \propto$
$t^0$
$\sqrt{t}$
$t^1$
$t^2$
$t^5$
$t^7$
$u(t) \propto t^9$
0.70M
0.50M
0.30M
0.20M
0.10M
0.05M
Fits
Predictions1
Predictions1
Predictions2
Predictions3

**Figure S5. Upon diverse kinetics, predictions enable identification of a true model among models of increasing complexities.** (A-E) Five models, with increasing complexities from left to right, are compared on the quality of their fits and predictions under data that are simulated from the true model (M3). From M3 to M2 and to M1, at each step one regulation is removed to generate simpler models. These could represent mutants of M3 where the corresponding kinase activities are removed resulting in loss of feedbacks regulations. From M3 to M4 two extra regulations are added to generate a complex 4-node model. This could represent potential hidden (yet undiscovered) regulations. Finally, M5 is generated by adding a whole new signaling branch (consisting of a sensor node and three regulations) to M3. Second row show fits (reds) of each model comparing to the train data (gray). Under parameters constrained for each model from these fits, next rows show model predictions. Third row shows models' predictions (examples of P1 in blue) compared to their corresponding testdata1 (steps, gray), and in forth row for linear inputs (blue). Fifth row shows models predictions (examples of P2 in yellow) compared to testdata2 (gray). Sixth row shows models predictions (examples of P3 in green) compared to testdata3 (gray). Results are from 5 independent fits of each model similar to Figures 5A-5C.  Specifically, on the results in this Figure (that is same as Figure 5), for all models, each fit is continued for additional 20 hours using fminsearch followed by the optimization algorithm given in Figure S1F to ensure all the models have reached their best fits.

Figure S6

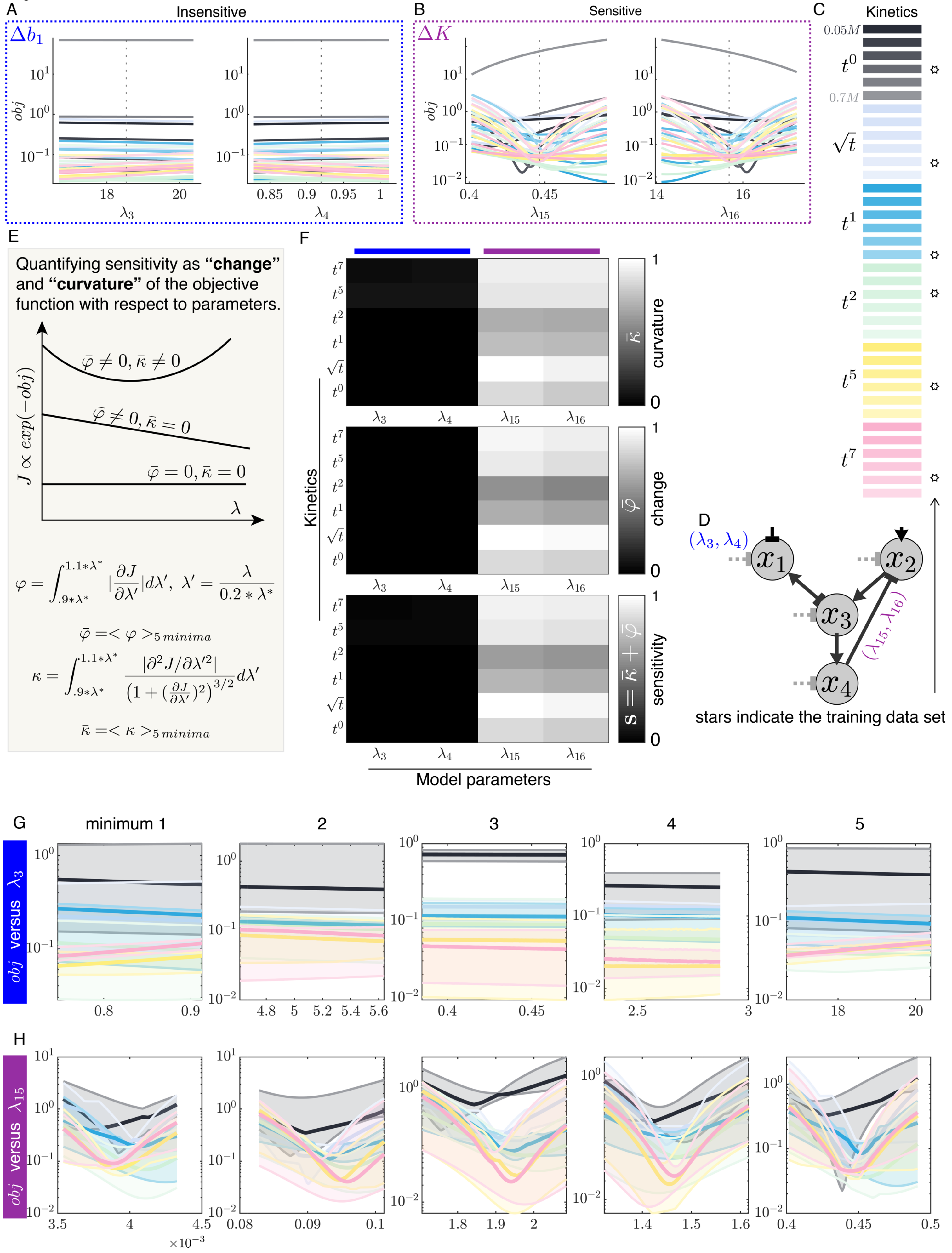
A
Insensitive
$\Delta b_1$
obj
$\lambda_3$
$\lambda_4$
B
Sensitive
$\Delta K$
$\lambda_{15}$
$\lambda_{16}$
C
Kinetics
$0.05M$
$t^0$
$0.7M$
$\sqrt{t}$
$t^1$
$t^2$
$t^5$
$t^7$
D
$(\lambda_3, \lambda_4)$
$(\lambda_{15}, \lambda_{16})$
$x_1$
$x_2$
$x_3$
$x_4$
stars indicate the training data set
E
Quantifying sensitivity as **“change”** and **“curvature”** of the objective function with respect to parameters.
$J \propto exp(-obj)$
$\bar{\varphi} \neq 0, \bar{\kappa} \neq 0$
$\bar{\varphi} \neq 0, \bar{\kappa} = 0$
$\bar{\varphi} = 0, \bar{\kappa} = 0$
$\lambda$
$\varphi = \int_{.9*\lambda^*}^{1.1*\lambda^*} |\frac{\partial J}{\partial \lambda'}| d\lambda', \ \lambda' = \frac{\lambda}{0.2*\lambda^*}$
$\bar{\varphi} = <\varphi>_{5\,minima}$
$\kappa = \int_{.9*\lambda^*}^{1.1*\lambda^*} \frac{|\partial^2 J/\partial \lambda'^2|}{(1+(\frac{\partial J}{\partial \lambda'})^2)^{3/2}} d\lambda'$
$\bar{\kappa} = <\kappa>_{5\,minima}$
F
Kinetics
curvature $\bar{\kappa}$
change $\bar{\varphi}$
sensitivity $S = \bar{\kappa} + \bar{\varphi}$
Model parameters
G
minimum 1
2
3
4
5
$obj$ versus $\lambda_3$
H
$obj$ versus $\lambda_{15}$
$\times 10^{-3}$

**Figure S6. Sensitivity analysis of parameters of the WT model trained upon diverse kinetics allows to screen for mutant responses**. (A-D) Sensitivity analysis of WT model with respect to parameters categorizes model parameters into two main groups of insensitive (e.g., $\lambda_3$ and $\lambda_4$ in A) or sensitive ($\lambda_{15}$ and $\lambda_{16}$ in B). In (A and B), the vertical black dotted line indicates the fit value of the corresponding parameter ($\lambda$*), and objective function (Figure S1F) is calculated around $\lambda$* upon all kinetic inputs (C). The 5 independent fits are used from Figures 5B and 5G where WT model (D) is trained with six kinetic input data (indicated with stars in C). (E) Sensitivity ($s$), defined as the sum of curvature ($\kappa$) and change (derivative, $\varphi$) of the objective function with respective to the parameter each integrated around $\lambda$* within 20%. Sensitivity for each model parameter is quantified over 5 independent fits (E) upon all kinetic inputs. It is then marginalized (summed over all final concentrations of each kinetic type) and normalized to the largest sensitivity. (F) For representative parameters, $\lambda_3$ and $\lambda_4$ (corresponding to **Δ**$b_1$, blue) and $\lambda_{15}$ and $\lambda_{16}$ (corresponding to **Δ**K, purple), sensitivity analysis predicts insensitive and sensitive mutants, respectfully. (G-H) An analysis of sensitivity over 5 independent fits indicate that the results are independent of the fits. (G) An example of insensitive parameter ($\lambda_3$) preserves this characteristic over all minima while $\lambda_3$* are different across them. (H) Similar result for sensitive parameters ($\lambda_{15}$). In (G-H), thick lines are median and shaded area are the interquartile range of all final concentrations for each kinetic type. Colors are according to the bar in (C). This result is consistent for all model parameters across independent fits.

Figure S7

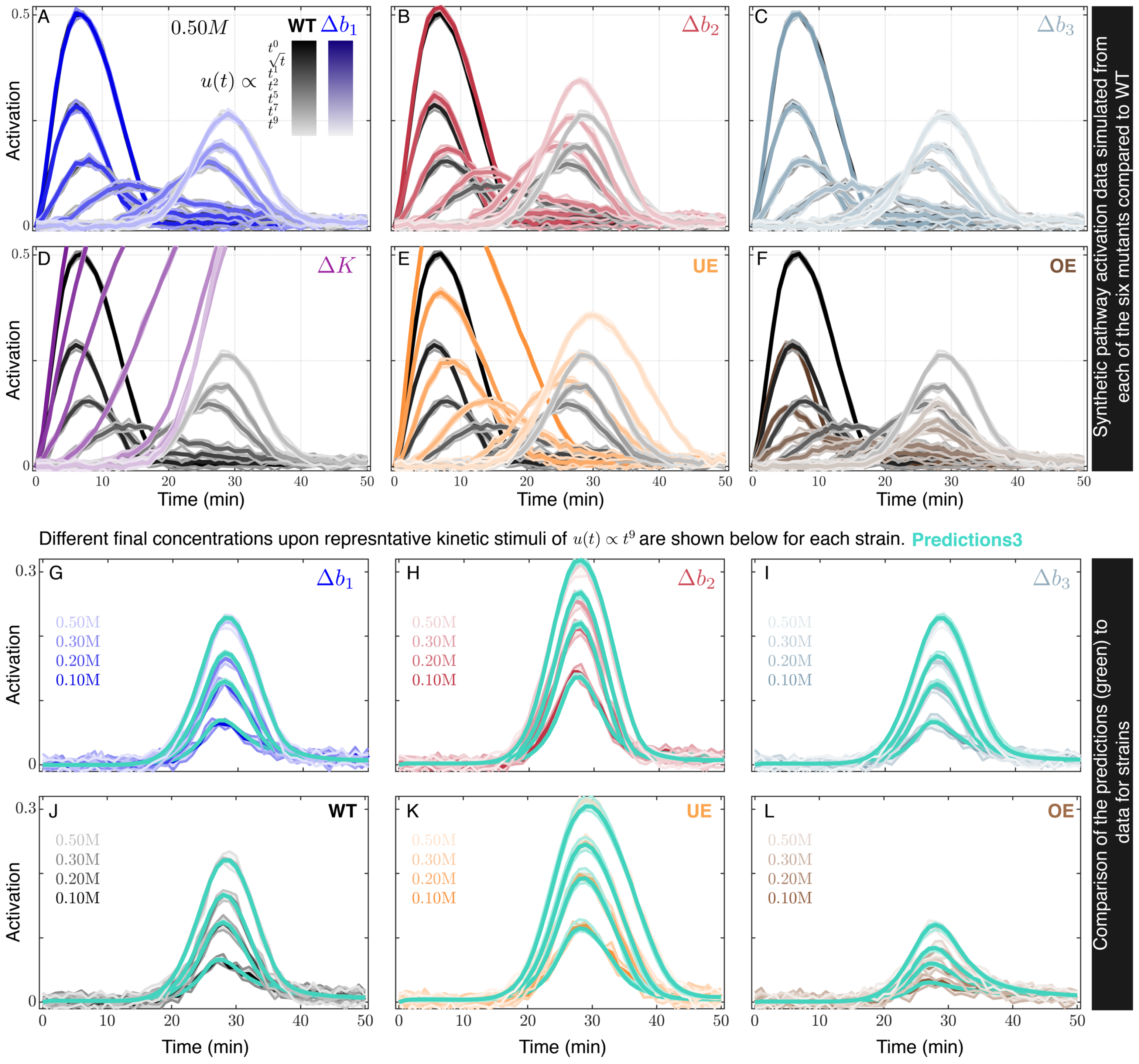

A
0.50M
WT Δb1
u(t) ∝
t^0
√t
t^1
t^2
t^5
t^7
t^9
B
Δb2
C
Δb3
D
ΔK
E
UE
F
OE
Activation
Time (min)
Synthetic pathway activation data simulated from each of the six mutants compared to WT
Different final concentrations upon represntative kinetic stimuli of u(t) ∝ t^9 are shown below for each strain.
Predictions3
G
H
I
J
K
L
WT
0.50M
0.30M
0.20M
0.10M
Comparison of the predictions (green) to data for strains

**Figure S7. Kinetically diverse stimuli enable to predict mutants' response dynamics.** (A-F) Synthetic signaling data simulated from mutated models upon kinetic inputs. Upon diverse kinetics (inset in A) and using parameter set $\boldsymbol{\Lambda}^0$ (Table S1), signaling activations are simulated from six mutants (given in Figure 6A) and each are compared to activation dynamics of WT strain (in gray) (STAR Methods). Some mutations are insensitive that's indistinguishable from the WT (A,C, $\boldsymbol{\Delta}b_1$ and $\boldsymbol{\Delta}b_3$) and others are sensitive (B,D-F, $\boldsymbol{\Delta}b_2$, **Δ**K, UE, and OE). Sensitivity analysis of WT model with respect to its parameters predicted insensitive (e.g., $\boldsymbol{\Delta}b_1$) versus sensitive (e.g., **Δ**K) mutants (Figure 6S). (G-L) Training the WT model on its responses to diverse kinetics enable to accurately predict its mutants' responses. Example pathway activation predictions (predictions3 in green) are compared to their corresponding synthetic data for $\boldsymbol{\Delta}b_1$ (G, blue), $\boldsymbol{\Delta}b_2$ (H, red), $\boldsymbol{\Delta}b_1$ (I, teal), WT (J, gray), $\mathrm{UE}$ (K, orange), $\mathrm{OE}$ (L, brown) under representative $t^9$ kinetic inputs. Mutant predictions (green) are generated under $\boldsymbol{\Lambda}$* (5 independent best parameter sets constrained by fitting WT model to its kinetically diverse data, Figures 5B and 5G) while mutants synthetic data (blue, red, teal, orange, brown) are simulated under $\boldsymbol{\Lambda}^0$ (Table S1 and Methods). Same comparison for the mutant **Δ**K is given in Figure 7D. Quantification of prediction (P1, P2, P3) errors over all 54 kinetics (Figure 1H) for each of the six mutants is given in Figure 7E.

**Table S1. Constrained parameter set that was used to simulate all the synthetic data in this paper from WT or mutant models.**

| | | | | |
|---|---|---|---|---|
| $\lambda_1 = 76.866$ | $\lambda_6 = 1000.000$ | $\lambda_{11} = 0.018$ | $\lambda_{16} = 7.639$ | $\lambda_{21} = 0.016$ |
| $\lambda_2 = 4.243$ | $\lambda_7 = 5.075$ | $\lambda_{12} = 0.071$ | $\lambda_{17} = 0.019$ | $\lambda_{22} = 1.170$ |
| $\lambda_3 = 0.093$ | $\lambda_8 = 1000.000$ | $\lambda_{13} = 136.332$ | $\lambda_{18} = 0.001$ | |
| $\lambda_4 = 941.483$ | $\lambda_9 = 0.001$ | $\lambda_{14} = 62.558$ | $\lambda_{19} = 1.018$ | |
| $\lambda_5 = 0.001$ | $\lambda_{10} = 456.755$ | $\lambda_{15} = 0.200$ | $\lambda_{20} = 4.369$ | |

These parameters are resulted from fitting the WT model shown in Figure S1K to Hog1 nuclear localization data of two steps of 0.2M and 0.4M simultaneously (Figure S1M). Among 10 fits shown in Figure S1M**,** this parameter set is chosen based on the quality of the predictions of model for pathway activation dynamics.
